# Mixed selectivity coding of content-temporal detail by dorsomedial posterior parietal neurons

**DOI:** 10.1101/2022.07.16.500237

**Authors:** Lei Wang, Xufeng Zhou, Fu Zeng, Mingfeng Cao, Shuzhen Zuo, Jie Yang, Makoto Kusunoki, Huimin Wang, Yong-di Zhou, Aihua Chen, Sze Chai Kwok

## Abstract

The dorsomedial posterior parietal cortex is part of a higher-cognition network implicated in elaborate processes underpinning memory formation, recollection, episodes reconstruction, and temporal information processing. Neural coding for complex episodic processing is however under-documented. Here we revealed a set of neural codes of ‘neuroethogram’ in the primate parietal cortex. Analyzing neural responses in macaque dmPPC to naturalistic videos, we discovered several groups of neurons that are sensitive to different categories of ethogram-items and to low-level sensory features, and saccadic eye movement. We also discovered that the processing of category and feature information by these neurons is sustained by accumulation of temporal information over a long timescale up to 30 s, corroborating its reported long temporal receptive windows. We performed an additional behavioral experiment and found that saccade-related activities could not account for the mixed neuronal responses elicited by the video stimuli. We further observed monkeys’ scan-paths and gaze consistency are modulated by video content. Taken altogether, these neural findings explain how dorsomedial PPC weaves fabrics of ongoing experiences together in real-time. The high dimensionality of neural representations should motivate us to shift the focus of attention from pure selectivity neurons to mixed selectivity neurons, especially in increasingly complex naturalistic task designs.

**HIGHLIGHTS:**

- Neural codes for “neuroethogram” in macaque dorsomedial parietal cortex
- Parietal neural codes exhibit mixed selectivity of event features
- Dorsomedial PPC neurons support a long temporal receptive window for episodes
- Saccadic movement could not explain away mixed neuronal responses
- Consistency in scan-path and gaze shown across viewing repetitions

## INTRODUCTION

In an ever-changing environment, massive amounts of multidimensional information embedded in continuous events rushes into the cognitive system. The neural system has to extract pieces of meaningful information, integrate, and encode them into memory systems in time for future needs. The dorsomedial posterior parietal cortex (dmPPC), which consists of the paracentral Area 7 and precuneus (Cavanna & Trimble, 2006), is part of the posterior-medial memory system (Ranganath & Ritchey, 2012) and has strong and wide-spread anatomical connections with its adjacent structures, including the early visual cortex, sensorimotor regions, medial temporal areas, and prefrontal cortex in both macaque monkeys and humans (Cavanna & Trimble, 2006; Kravitz et al., 2011; Morecraft et al., 2004).

A wealth of studies demonstrated that the dmPPC plays critical roles in multifaceted cognitive processes, including visual-spatial attention and locomotion processes in egocentric environment (Bartels et al., 2008; Ghaem et al., 1997), sensorimotor transformation processes, such as object manipulation (Gardner et al., 2007), execution and observation of reaching-to-grasp behaviors (Diomedi et al., 2020; Evangeliou et al., 2009), as well as representations of enumeration (Harvey et al., 2013), self-related processing (Cavanna & Trimble, 2006), and episodic memory formation and retrieval (Brodt et al., 2018; Brodt et al., 2016). The dmPPC is part of an integral hub for extracting and scaffolding information in real-time from the environment (Reagh & Ranganath, 2021) and with other agents (Freedman & Ibos, 2018; Kravitz et al., 2011).

Given the region’s myriad functions, the conventional logic of stimulus-response models might not be adequate for studying neuronal responses to complex stimuli and their interactions. To take an analogy, neurons in high-order brain areas such as the prefrontal cortex (Rigotti et al., 2013) and the parietal cortex could show mixed selectivity properties to different stimuli (Fusi et al., 2016; Wallach et al., 2021) and processes such as decision-making (Erlich et al., 2015) and visuomotor coordination (Diomedi et al., 2020). Indeed, Platt and his colleagues recently showed that neurons in PFC and OFC were engaged in valuing social information with a measure known as “neuroethogram” (Adams et al., 2021). A neuroethogram is defined as fitting neural activities to ethograms, which is a technique for annotating species-typical behavior. Considering that the primate Area 7 is part of a social interaction network (Sliwa & Freiwald, 2017) and a posterior-medial memory network (Ranganath & Ritchey, 2012), we predict that dmPPC neurons process information embedded within complex behaviorally meaningful events. Since the capacity of single neurons to integrate multiple variables flexibly should enhance the organism’s ability in processing nonlinear integration of multiple information sources (Vaccari et al., 2022), we reasoned that this capacity should be especially important for dealing with complex episodic information like video content.

In addition to leveraging on multifunctional features contained in naturalistic videos, we were mindful that temporal information is another fundamental aspect of events (Clewett et al., 2019). Since information is carried out over distinct timescales, we know that the ability of information accumulation changes from the primary sensory cortex to the high-order cortex (Hasson et al., 2015; Hasson et al., 2008). These studies propose that brain areas are organized with hierarchical time receptive windows (TRW) so that brain regions with long TRW will accumulate transient sensory signals from short TRW brain areas for processing. The human precuneus plays an essential role in temporal information integration of movies of up to 12 s (Andric et al., 2016; Hasson et al., 2008). It is unclear how neurons in the dmPPC might dynamically assemble such temporal details to support such episode processing.

To address these two issues on processing multiplex content and passage of time, we combined an ethogram methodology with dynamic cinematic material, together with multi-unit extracellular electrophysiology in awake macaque monkeys, to elucidate how dmPPC neurons process mixed selectivity representation of naturalistic content over time. In an additional eye movement experiment we found that gaze behaviors and saccade-related activities could not fully account for the mixed neuronal responses elicited by the video stimuli.

## RESULTS

In this study, we used five macaque monkeys. To study mixed selectivity coding, we had 3 monkeys view 18 different movies (categorically Primate/Non-Primate/Scenery content-type) while we performed extracellular action potential recording targeting the dorsomedial posterior parietal cortex (with one of them having eye movement simultaneously recorded). On each day, the monkeys watched 3 different movies, each for 30 repetitions. These movies were custom edited to contain both content and temporal information (Figure 1A; Table S1). In total, we recorded extracellular activities of 375 units (monkey Jupiter: 164; monkey Mercury: 157; monkey Galen: 54) (Figure 1B, Figure S1; Methods; Movie S1). In addition, we performed an eye movement experiment on two further monkeys to examine saccadic events and gaze consistency during viewing.

**Figure 1.**
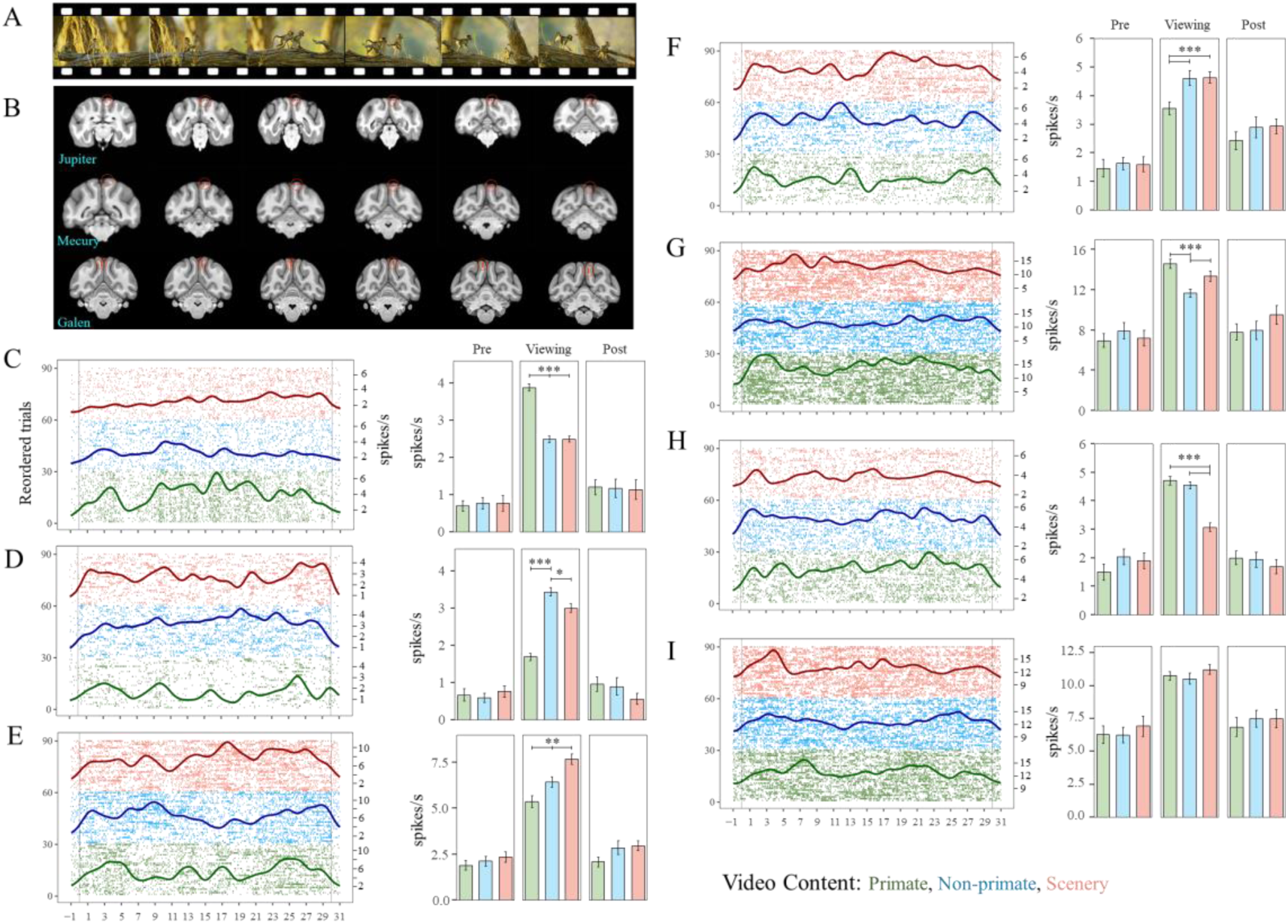
Experimental procedure, recording sites, and neuron classification. (A) Example video (a Primate-video) used in the study. Each day, the monkeys watched three different 30-s videos, each for 30 repetitions in 6 blocks; (B) Reconstruction of recording sites (circled in red) overlaid on T1 images; (C-I) Raster plots (left panels) of reordered trials (left axes) overlaid by spike-density histograms with 100 ms Gaussian kernel smoothing (right axes) and firing rate comparisons (right panels) of 7 representative neurons responding to different video content-types. All show a significantly higher firing rate during video viewing than pre/before and post/after video presentation (Ps < 0.05). In raster plots, the X-axis indicates the time course of the video, and vertical lines represent the onset or offset of the video display; each row is associated with a trial. Trials are re-ranked by video content-types (yellow: Scenery, blue: Non-primate, green: Primate). Three example content-sensitive neurons showed significantly higher firing rates to primate (C:#PC0056, Primate), non-primate (D:#PC0040, Non-primate), and scenery (E:#PC0114, Scenery) content-type. Firing rates of three content-sensitive neurons exhibited the lowest activity to primate (F:#PC0232, Non-primate-Scenery), non-primate (G:#PC0205, Primate-Scenery), and scenery (H:#PC0249, Primate-Non-primate) content-types. (I) A Content Insensitive example neuron (#PC0192) exhibited equal firing rates across different video content-types. Error bars: SEM. * P < 0.05, ** P < 0.01, *** P < 0.001.

### Classification of neurons by their specificity to video’s content-type

By comparing the neural spiking rates to each of the three video content types, we observed that 33.07% (124/375) of the neurons exhibited significant (one-way ANOVA, Ps < 0.05) content-sensitive activity in the videos (Figure 1C-H), while 66.93% (251/375) showed no difference on firing rate across video contents (content-insensitive, Figure 1I). Among these content-sensitive neurons, 52.4% (65/124) of the units had a higher mean firing rate for Primate videos (Primate), whereas 8.9% (11/124) and 16.9% (21/124) had a higher mean firing rate to Non-primate (Non-primate) and Scenery (Scenery) videos, respectively (Figure 1C-E; segments of frames and raster plot of spikes for a Primate video in Figure S2A-B). In contrast, 10.5% (13/124) of these content-sensitive neurons discharged less to Primate videos (Non-primate-Scenery, Figure 1F), and 3.2% (4/124) and 8.1% (10/124) discharged less to Non-primate (Primate-Scenery, Figure 1G) and Scenery contents (Primate-Non-primate, Figure 1H). We found that the dorsomedial parietal neurons respond differentially to different video content-types, with some of them to the primate content, which is consistent with previous findings that a portion of this part of the monkey medial parietal cortex is activated by social interaction of conspecifics (Sliwa & Freiwald, 2017). We also examined whether and to what extent the temporal spiking patterns during viewing can differentiate the representations of categorical content. We trained a linear multiclass support vector machine (SVM) classifier with firing rate with 1-s time bins using a leave-one-out cross-validation approach for each neuron (see Methods Video Content Type Discriminability). 40.80% (153/375) of the neurons exhibited a significant decoding ability when compared to a label-shuffled permutation statistical threshold (Ps < 0.05).

### Multiplex representation of ethogram-items and low-level features in dmPPC neurons

We analyzed the videos in detail by employing a semi-automatic frame-by-frame annotation of ethogram schema, which contains a subset of binary time series of observable social and non-social events (Adams et al., 2021). To investigate how individual neuron encodes the dimensions of informative dynamic natural context, we fit a LASSO elastic network regulation regression for each neuron to fit the averaged single neural activities with the binary labeling of the 52 ethogram-items (Table S2) and four low-level visual features (Figure S3). The analysis produced a collection of non-zero coefficients. As shown with an example neuron, the model with the lowest mean squared error would be chosen (Figure S4A-C). We validated the chosen model by demonstrating a significant relationship with the predicted neural firing rates (F(1, 450) = 121.5, R^2^ = 0.213, P < 10^-5^; Figure S4D). By applying this feature selection procedure for all neurons, our model indicated that the activity of a large percentage of neurons would either positively or negatively be influenced by a number of ethogram items depicted in the videos (range from 3.73% to 74.13%; Figure 2A-B).

**Figure 2.**
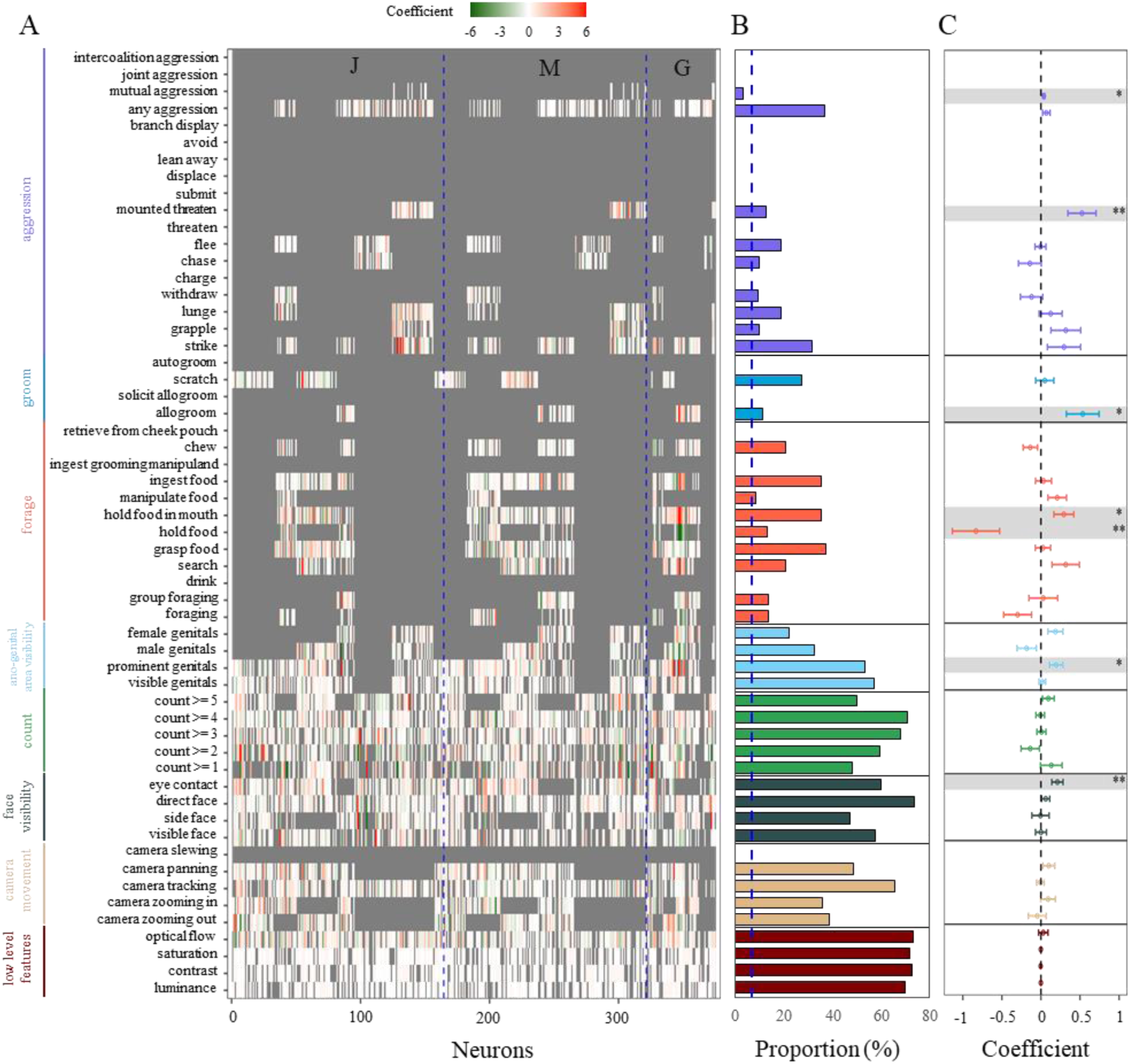
dmPPC neurons respond to social and non-social events in videos. (A) Effects of neuronal responses to 4 low level features (dark red) and 52 ethogram-items (seven categories of the ethogram organized by 7 different colors) obtained by a least absolute shrinkage and selection operator (LASSO) regression analysis. Each row stands for an item out of all 56 items, while each column refers to one neuron. Blue dashed lines showed neurons acquired from three different monkeys (J = Jupiter, M = Mercury, G = Galen). (B) Proportion of neurons responsive to each item. The blue dashed line indicates chance level. (C) LASSO coefficient for each item tested against zero. Selected features and its modulation values of an example neuron were shown in Figure S2C. Error bars: SEM over neurons. * P < 0.05, ** P < 0.01.

A large proportion of units respond to facial (visible face: 57.87%; side face: 47.47%; direct face: 74.13%; eye contact: 60.27%) and genital (visible genitals: 57.60%; prominent genitals: 53.87%; male genitals: 33.07%; female genitals: 22.4%) features (Figure 2A-B). With prominent ethogram-items, we ran one-sample t-tests to test for their modulatory effects. The results revealed that eye contact (t(225) = 2.921, P = 0.004, Cohen’s d = 0.194), prominent genitals (t(201) = 2.249, P = 0.026, Cohen’s d = 0.158), holding food in mouth (t(132) = 2.298, P = 0.023, Cohen’s d = 0.199), allogroom (t(43) = 2.561, P = 0.014, Cohen’s d = 0.386), mounted threaten (t(48) = 2.911, P = 0.005, Cohen’s d = 0.416), and mutual aggression (t(13) = 2.643, P = 0.020, Cohen’s d = 0.706) significantly enhanced firing rates, whereas holding food (t(50) = -2.750, P = 0.008, Cohen’s d = 0.385) reduced neuronal activities (Figure 2C). The proportion of the neurons modulated by each of the features are higher than chance (see Methods Chi-squared Simulation) (Figure 2B).

We checked whether dmPPC neurons would play a role in the processing of low-level features (cf. monkey early visual cortex in (Russ & Leopold, 2015)). In the same LASSO model, low-level features tuned a large proportion of the dmPPC neuronal responses (“low level features” in Figure 2A & dark red bars in 2B). A large subset of neurons was tuned by luminance (70.40%, 264/375), contrast (73.07%, 274/375), saturation (72.00%, 270/375), and optical flow (73.60%, 276/375) respectively. To elucidate the separate contributions of high-vs. low-level features, we performed a separate cross-validation LASSO regression model to identify how low-level features contribute to the neural activities. Taking the neuron #PC0087 as an example, compared to our original full model (see Figure S4D), the R^2^, or the explanatory power of the regression model, remained statistically significant (F(1, 450) = 78.85, R^2^ = 0.147, slope = 0.156) once the low-level feature was removed. This implies that we would not be able to exhaustively capture all the variations contributed by all low-level features (e.g., local spectral and spatiotemporal elements) to reach a 100% fitting of the neural activities.

Previous studies suggested that the neuronal latencies of responses to the onset of stimuli in the posterior parietal cortex range from 45.2 ms in LIP (Bisley et al., 2004) to 98 ms in Area 7a (Barash et al., 1991; Bushnell et al., 1981). To test whether the latency had any effect on our results, we realigned the spike train to the onset of the video’s presentation with either a 40-ms or a 100-ms latencies to fit the LASSO feature selection algorithms. By comparing these with our original results, the proportions of most selected features were not affected by the latencies realignments in both cases (Ps > 0.223), except that more neurons respond to male genitals with 100 ms realignment (0 ms vs.100 ms: χ^2^(1) = 32.988, P < 10^-5^; 40 ms vs. 100 ms: χ^2^(1) = 38.429, P < 10^-5^) but fewer neurons were modulated by the feature of flee (0 ms vs. 40 ms: χ^2^(1) = 15.991, P < 0.001; 0 ms vs. 100 ms: χ^2^(1) = 25.126, P < 10^-5^) and any aggression (0 ms vs. 40 ms: χ^2^(1) = 10.882, P < 0.001; 0 ms vs. 100 ms: χ^2^(1) = 18.961, P < 0.001) in both latencies (Figure S5A-B). Yet, the coefficients of these selected features showed no differences against 0 ms latency alignments in both 40 ms (Ps > 0.524) and 100 ms (Ps > 0.362) realignments. Since these control analyses indicated that shifting the latency by 1 (40 ms) or 2 (∼100 ms) frames did not change our main results, we have analyzed the data with no latency-shifting for the remaining analyses.

It has been reported that neural activity in PPC is involved in primate saccadic behavior (Andersen et al., 1990). To evaluate the extent to which the neural activation was modulated by saccades during video-viewing, we repeated the LASSO feature selection algorithm but now taking saccades as the 57^th^ feature along with all other ethogram items on trial-by-trial basis (see Methods Eye Movement Analysis) for data acquired from monkey Galen. The results demonstrated that 85.19% (46/54) neural activities were increased by saccadic behaviors (t(45) = 3.271, P = 0.002). At the group level, however, the proportions (Ps > 0.541) and coefficients (Ps > 0.560) of neurons responding to each of the 56 key items showed no significant differences between whether saccadic data was included as a feature (Figure S6). This suggests that even when potential contributions by eye movements were regressed out, the neuronal selectivity responses to the key features and visual content remain.

To illustrate the multiplex nature, we then performed an intersection analysis (Bastian et al., 2009) and found that almost all (94.4%; 354/375) units showed mixed selectivity representations to at least three ethogram categories, 1.6% (6/375) neurons modulated by the combination of two ethogram categories, and only 4% (15/375) units selectively respond to one single ethogram category (Figure 3 & Movie S2).

**Figure 3.**
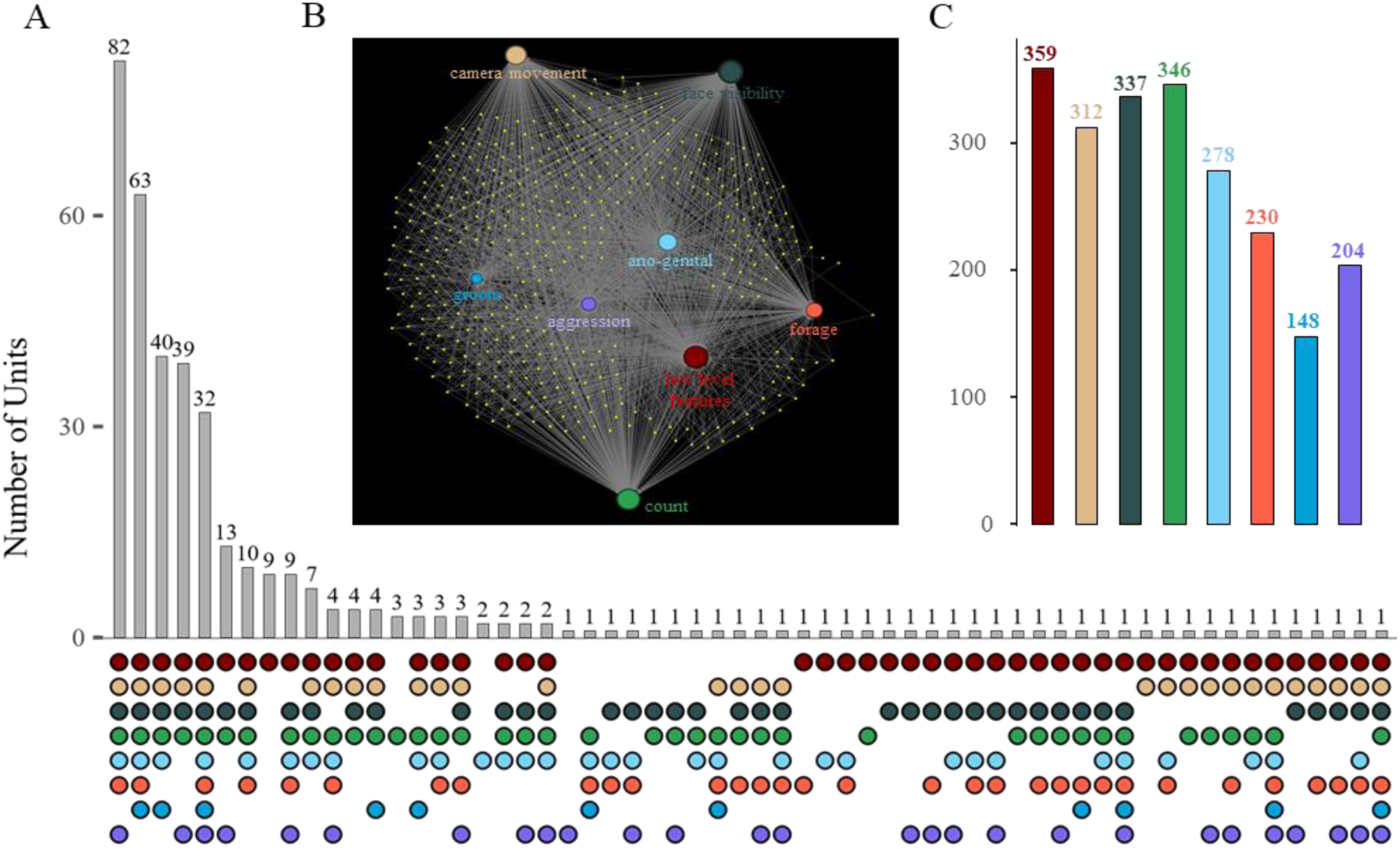
dmPPC neurons demonstrate mixed selectivity representations. (A) Distribution of neurons and their composition for mixed selectivity representations. Gray bars show the numbers of units exclusively modulated by combinations of mixed ethogram features, with their composition shown in the bottom panel. Color coding here is the same as Figure 2 and Figure 3B. (B) Demonstration for dmPPC cell ensembles for their mixed selectivity coding. Each small yellow dot denotes a neuron. The eight circles with labels refer to the eight feature categories (low-level features and 7 ethogram categories), with their size proportional to the number of neurons modulated by that category. The connecting lines refer to the relationship between neurons and feature categories. See also an interactive illustration of the multiplex behavior of individual neurons on Movie S2. (C) Number of neurons that responded to each of the ethogram categories. For example, the category “camera movement” including multiple camera motions modulate the discharge of about 83.20% (312/375) percentage of all units. The category “count”, that is the number of animals visible, influenced a significant portion of all units (92.27% (346/375).

### Long temporal receptive window sustained by dmPPC neurons

In light of the proposal that the parietal association cortex exhibits information accumulation over long timescale (Honey et al., 2012; Murray et al., 2014; Runyan et al., 2017), we hypothesized that the dmPPC cells might help scaffold the dynamic events temporally. To test the temporal accumulation hypothesis, we constructed a multiclass SVM classifier with stepwise accumulated sequential spiking using 1-s time bins across the videos (light green dots/line in Figure 4A). For this example neuron (*#PC0361*), we showed that accumulated 1-s epoch decoding produced a significantly better decoding performance than shuffled data (t(29) = 11.013, P < 0.001, Cohen’s d = 2.011; dark green dashed line in Figure 4A), with prediction accuracy increased as a function of accumulated time points (R^2^ = 0.757, P < 10^-5^).

**Figure 4.**
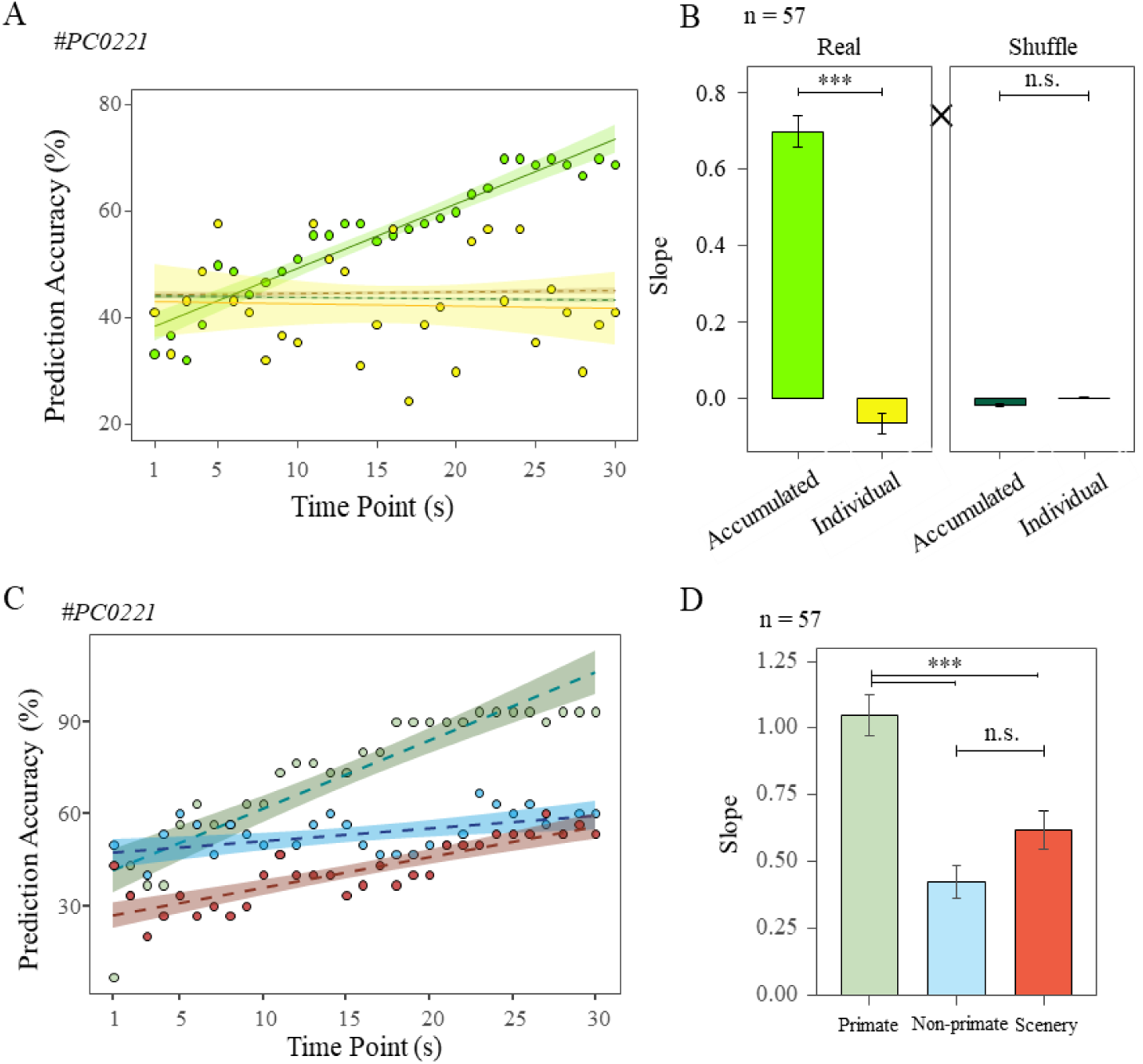
dmPPC neurons accumulate temporal information with long temporal receptive windows. (A) The decoding performance of the example neuron (*#PC0221*) positively correlates with cumulative spiking sequences (light green) but not with momentary neural activity (yellow). We used two sets of SVM decoding exercises to verify this property. First, we used cumulative spikes in 1-s time bins for 1st to 30th timepoint (accumulated sequence; light green) and compared it to the significant statistical threshold (dark green). Second, we used spikes in each individual time-point (yellow) and compared it with the permuted significant statistical threshold (dark red). The four lines represent linear regression for these four SVMs for an example neuron. The dots refer to decoding performance of each timepoint for cumulative and momentary conditions. (B) A Sequence (Accumulative/Individual) × Approach (Real/Shuffle) two-way ANOVA revealed that the mean slope of population neurons (n = 57) was higher for real and cumulative sequences than for both shuffled control data and individual 1-s time binned spike data (Ps < 0.001). (C) The decoding performance for each video content positively correlates with cumulative spiking sequences. (D) One-way ANOVA and post hoc analysis revealed that neurons in dmPPC had fast accumulation speed for primate video contents. **×** indicates the Sequence by Approach two-way interaction. Error bars: SEM across neurons. *** P < 0.001. ns: not significant.

For statistical inference, an identical SVM decoding procedure was applied but using neural activity for each time bin independently (yellow dots/line in Figure 4A; see Figure S8 for illustration). The prediction accuracy was not better than the permuted threshold and its slope was not different from zero (R^2^ = 0.006, P = 0.689; brown dashed line in Figure 4A). Planned paired t-tests confirmed that evidence accumulation is inherent in the temporal sequences rather than the single moments during which neurons fire (t(29) = 12.154, P < 0.001, Cohen’s d = 2.219).

Using this approach, 57 of the neurons showed a statistically significant pattern for information accumulation. To assess the strength of the information accumulation, we compared the slopes by crossing two factors, Sequence (Accumulative/Individual) × Approach (Real/Shuffle) and found a two-way interaction (F(1,56) = 339.307, P < 10^-5^, η^2^ = 0.531). These effects were derived from the stronger effects in accumulated real firing sequences than momentary neural firing (Real-Accumulated vs. Real-Individual: t = 25.603, P < 10^-5^, Cohen’s D= 2.919; left panel in Figure 4B) and no differences between the two shuffled conditions point (t = 0.645, P = 0.520; right panel in Figure 4B). Additional analysis revealed that these accumulation neurons had faster speed and stronger magnitude to accumulate conspecific relevant information than Non-primate and Scenery information (F(2,112) = 21.056, P < 0.001, η^2^ = 0.273; Primate vs. Non-primate: t = 6.337, P < 0.001, Cohen’s D = 1.183; Primate vs. Scenery: t = 4.378, P < 0.001, Cohen’s D = 0.817; Figure 4C-D). These findings show that dmPPC neurons accumulate information of dynamic events in an additive manner over the course of video viewing, especially for conspecific related events.

As a control analysis, we tested whether the effect was an artifact of growing firing sequences. We repeated the multiclass SVM decoding approach but now with smaller stepwise accumulated 40-ms time bins spiking sequences across the videos (in place of 1-s time bins). 31.37% (48/153) of the neurons sustained significant information accumulation, which is not statistically different from the performance with 1-s time bins (χ^2^(1) = 0.9279, P = 0.335, Figure S8), indicating that the dmPPC neurons exhibit its accumulation property irrespective to the length of the spiking sequences.

### Scan-paths over repetitions and gaze consistency modulated by video features

In the final set of analyses, we used data from an eye movement experiment and analyzed the gaze behavior of three monkeys (one of them had also participated in the electrophysiology experiment). Previous studies reported that non-human primates made anticipatory saccades to memorized events (Kano & Hirata, 2015), implying that primates exhibit consistency in gaze trajectory across repeated viewings. To test this, we calculated the pairwise scan-paths correlations across viewing repetitions and showed that monkeys exhibited significantly stronger similarity across viewing scan-paths for videos with conspecific activities (Primate vs. Non-primate and Primate vs. Scenery: *P*s < 0.05; Figure 5 A-B). We then ran a GLM regression to assess the change in scan-paths similarity as a function of repetition lag, which is measured by the time between any two paired viewings of the same video. We showed a lag effect that scan-paths similarity decreases with increasing repetition lag, but consistently for all three content-types (Primate: R^2^ = 0.782, P < 10^-5^; Non-primate: R^2^ = 0.668, P < 10^-5^; Scenery: R^2^ = 0.530, P < 10^-5^; Figure 5 C).

**Figure 5.**
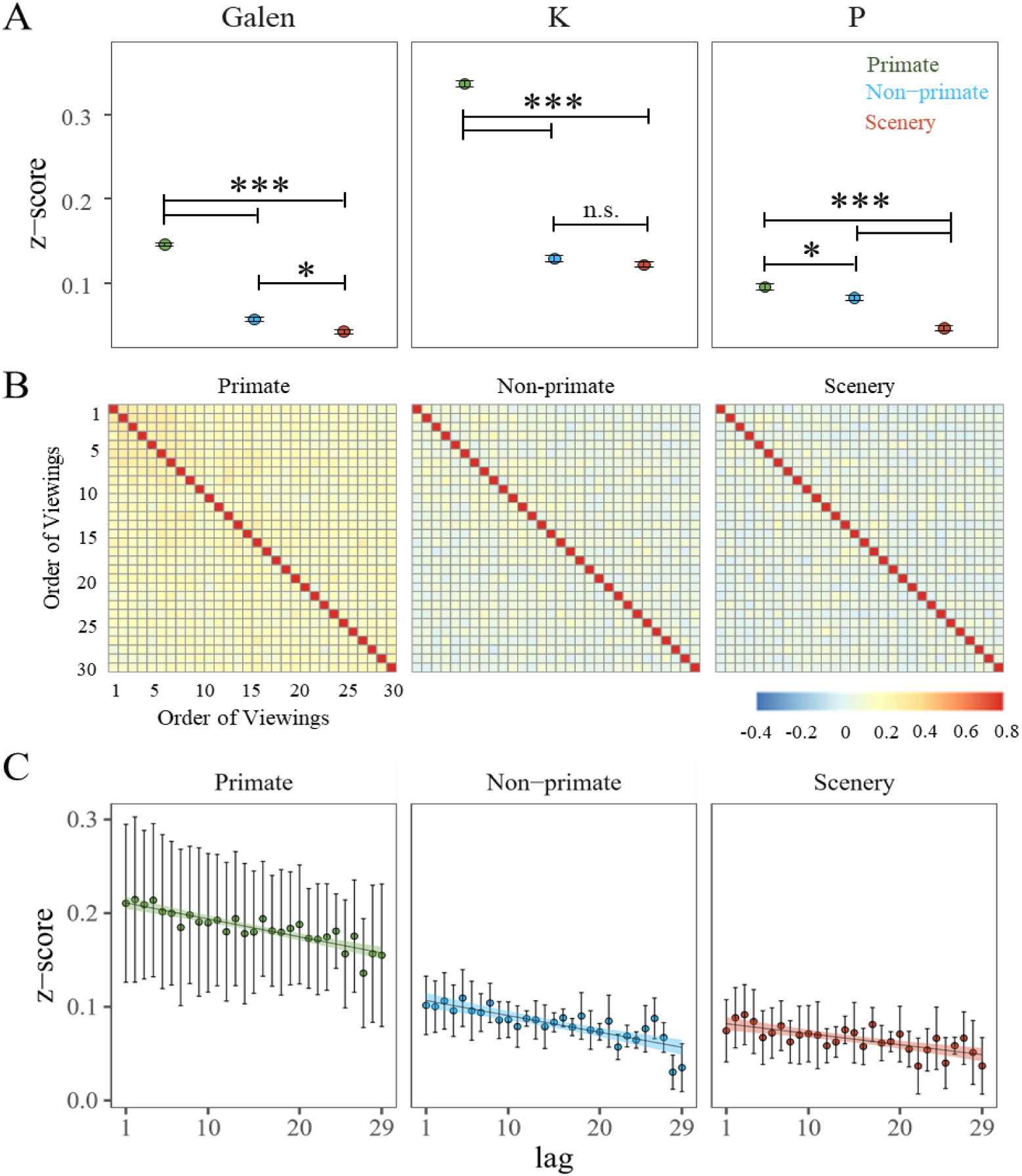
Scan-paths across viewings remain most steady for Primate video content-type. (A) Monkeys showed higher scan-paths similarities across viewing repetitions for Primate than for Non-primate and Scenery videos (Galen: F(2, 1302) = 588.808, P < 0.001, η^2^ = 0.475; t _Primate - Non-primate_ = 27.288, P < 0.001, Cohen’s D = 1.850; t _Primate - Scenery_ = 31.664, P < 0.001, Cohen’s D = 2.147; t _Non-primate - Scenery_ = 4.376, P < 0.001, Cohen’s D = 0.297; monkey K: F(2, 1302) = 1228.672, P < 0.001, η^2^ = 0.654; t _Primate - Non-primate_ = 42.212, P < 0.001, Cohen’s D = 2.862; t _Primate - Scenery_ = 43.615, P < 0.001, Cohen’s D = 2.957; t _Non-primate - Scenery_ = 1.403, P = 0.340, Cohen’s D = 0.095; monkey P: F(2, 1302) = 52.018, P < 0.001, η^2^ = 0.074; t _Primate - Non-primate_ = 2.620, P = 0.024, Cohen’s D = 0.178; t _Primate - Scenery_ = 9.847, P < 0.001, Cohen’s D = 0.688; t _Non-primate - Scenery_ = 7.227, P < 0.001, Cohen’s D = 0.490). Error bars: SEM across repetitions. (B) Heatmaps of averaged pairwise scan-path correlations of the three monkeys, showing significantly higher correlation for the Primate content-type eye data. (C) Scan-path similarity plotted as a function of repetition lag. Error bars in C refer to SEM across monkeys. ***: p < 0.001; *: p < 0.05; n.s.: No significant.

To identify the kind of features in the video that influence monkeys’ gaze consistency across viewings, we constructed a separate LASSO regression on the ethogram items using gaze consistency, that is a measurement of distribution of gaze central tendency on each frame across the 30 viewings. We obtained patterns of results that are in consistency with Adams et al. findings (Adams et al., 2021) that items such as animal count equal or more than 1, visible and prominent genital cues, hold food in mouth resulting in high gaze consistency, whereas items such as presentation of male genitals, manipulate food, allogroom on the screen leading to lower gaze consistency (Figure 6).

**Figure 6.**
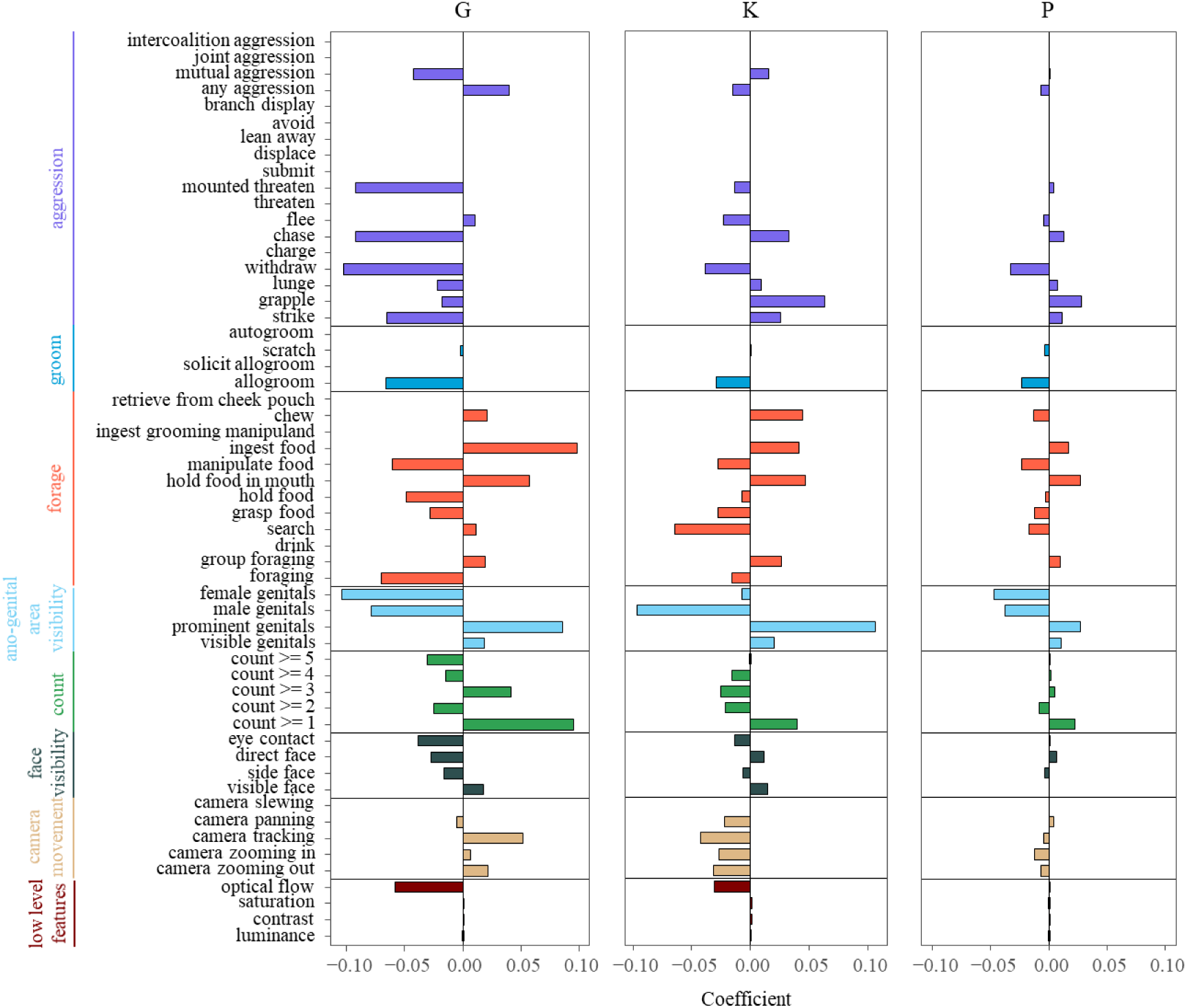
Monkeys’ gaze consistency modulated by the video features. Bars denote the non-zero coefficients chosen by the LASSO feature selection algorithm. Positive coefficients indicate items that result in high consistency across viewings whereas negative coefficients refer to low consistency across viewings. The 3 monkeys showed high agreement for a majority of items.

## DISCUSSION

Our findings revealed that neurons in the dorsomedial posterior parietal cortex (dmPPC) exhibit responses to an array of cinematic features. DmPPC neurons showed mixed selectivity responses to different categories of ethograms and low-level visual features. The amount of information embedded within neuronal spiking sequences were modulated by the convergence of multiple representations which in turn contribute to the read-out of different content-types. The processing of category and feature information by these neurons is sustained by accumulation of temporal information over a relatively long timescale.

According to dual-process models in social cognition, the medial posterior parietal cortex is part of the reflective system corresponding to a controlled social cognition processing (Lieberman, 2007; Satpute & Lieberman, 2006). Here, a large proportion of neurons respond to the presentation of foraging behaviors. This pattern might be related to evidence that neurons in macaque’s Area 7 express its intentions before actions (Snyder et al., 1997) and that Area 7 strongly responds to conspecific social interactions in comparison to non-social interactions by inanimate objects (Sliwa & Freiwald, 2017). Further to Klein’s observations that neuronal activity in the primate lateral intraparietal cortex signaled values of conspecific genital cues (Klein et al., 2008), such modulation is also observed in dmPPC neurons in the present study. Since neurons in dmPPC are consistently modulated by the observation of social interactions, such as inner group grooming behaviors (allogroom and scratch) and aggression (chase, strike, flee), our findings strengthen the notion that the posterior parietal cortex plays an interface role in the integration of multifaceted information for social cognition. All in all, the neurons in the dmPPC demonstrate mixed selectivity representation (Vaccari et al., 2022) to distinct as well as combinations of features (e.g., aggressive behavior and allogroom), implying a role of dmPPC in the computation for multiplex information in rich fast-changing environments (Fusi et al., 2016; Johnston et al., 2020; Murray et al., 2017). In several control analyses, we further confirmed saccadic movement could not explain away the mixed neuronal responses and that transmission latency also did not affect the overall main results.

Another key finding of our study is that the neurons in the monkey dmPPC accumulated representations across the duration of video to support the construction of episodes, providing evidence that the PPC sustains a long temporal receptive window. Indeed, the dmPPC has been proposed to code information with long timescales (Hasson et al., 2008; Runyan et al., 2017). A human fMRI study reported that medial posterior parietal cortex (precuneus) accumulates information of video stimuli up to 12 s (Hasson et al., 2008). We argued that the long TRW allows the dmPPC to accumulate the continuous information of multifaceted representations from unimodal or integration of cross-modal inputs (Gilissen & Arckens, 2021) to support the processing of streams of episodic information. This aligns with the higher temporal dynamics in the precuneus when remembering the unfolding of events that included a high density of experience units (Jeunehomme et al., 2022). The primate dmPPC has dense connections to the hippocampal formations in the primate (Kravitz et al., 2011). Given the known importance of schema cells in the primate hippocampus (Baraduc et al., 2019) and how such cells might code for information about space and nonspatial elements of the environment for both perceptual and mnemonic experiences (Gulli et al., 2020; Zhang et al., 2022), it is likely that the two structures support an information abstraction system that is driven by a broad range of behaviorally relevant inputs.

Functional MRI studies using dynamic movie stimuli reveal that the medial PPC is involved in the motion information processing in both humans and monkeys (Bartels et al., 2008; Russ & Leopold, 2015). This is in line with our observation that neural activity in dmPPC were modulated by motion features, including optical flow (Raffi & Siegel, 2007) and motions caused by camera movement and animal appearance (count), aggressive behaviors (e.g., chase, strike, flee), and other features correlated with the fluctuations in early visual areas (Russ & Leopold, 2015). Neuronal responses to the convergence of multiplex dimensional stimuli suggest that dmPPC is involved in the integration of inputs from inner projections in the PPC (Andersen et al., 1987) and from the early visual cortex (Robinson et al., 1978). Moreover, in the eye movement experiment, we further demonstrated that scan-paths are modulated by video content-type across repetitions. Analysis on gaze consistency across viewings also confirmed how specific ethogram items embedded in the videos could result in different viewing patterns (Adams et al., 2021).

Compared to static stimuli, the use of dynamic naturalistic videos, which meets the ethological validation from sensory to social cognition research (Adams et al., 2021; Mosher et al., 2014; Testard et al., 2021), helped us yield significant findings in the present study. We acknowledge that the analyses and their conclusions are based on correlational measures between action potentials and cognitive indices. In future work, micro-stimulation by electrical or targeted pharmacological intervention would be instrumental for a detailed elucidation of the complex cognitive roles carried out by the primate dorsomedial posterior parietal neurons.

## RESOURCE AVAILABILITY

### Lead contact

Further information and requests for resources should be directed to the Lead Contact, Sze Chai Kwok.

### Data and code availability

Raw electrophysiological data, analysis code, and processed data supporting the conclusions of this study are available upon request.

## EXPERIMENTAL MODEL AND SUBJECT DETAILS

### Subjects

Five male rhesus macaques (*Macaca mulatta*) (8.66 ±0.59kg) with a mean age of 6.6 years old served as subjects in this study. Among them, we recorded extracellular electrophysiological activities from two monkeys (monkey Jupiter: 6y, 8.3kg; monkey Mercury: 6y, 8.6kg) and both neural activities and eye movements from a third monkey (monkey Galen: 8y, 9.2kg). Furthermore, we performed an eye movement experiment on two additional monkeys (monkey K: 7y, 9.3kg; monkey P: 6y, 7.9kg; Table S2).

All monkeys were single-housed with a 12:12 (7: 00am/7:00 pm) light-dark circle and kept within the temperature range of 18 - 23℃ and humidity between 60% - 80%. The animals were fed twice a day with each portion of at least 180g monkey chow and pieces of apple (8:30 am/4:00 pm). Water was limited during recording days. All animal care, experimental, surgical procedures, and pre/post surgical care were approved by the Institutional Animal Care and Use Committee (permission code: M020150902 & M020150902-2018) at East China Normal University.

Before this study, monkey Jupiter and monkey Mercury participated in a temporal order judgment behavioral experiment (Wang et al., 2020; Zuo et al., 2020). Monkeys Galen, K and P were trained on fixation and saccadic tasks and participated in several oculomotor studies with neuronal activities recorded from right medial temporal and medial superior temporal areas (Jia et al., 2021).

## METHOD DETAILS

### Experimental procedure and overview

During this study, the monkeys sat in a custom-manufactured Plexiglas monkey chair (29.4 cm × 30.8 cm × 55 cm) with a head-fixed (see Surgery) in front of a 19-inch screen (An-190W01CM, Shenzhen Anmite Technology Co. Ltd, China) mounted on a stainless-steel platform. Monkeys’ eyes are about 60 cm and 62 cm away from the screen’s top edge and bottom edge, respectively. Water was delivered by a distributor (5-RLD-D1, Crist Instrument Co., Inc., U.S.) as a reward.

In each session, the monkeys watched 3 different 30-s video footage presented with PsychoPy (PsychoPy 3.1.2, PsychoPy), each for 30 repetitions arranged in 6 blocks. The same list was repeatedly presented in two consecutive days (12 days in total). In total, each video was watched 60 times. 1ml water was delivered at the beginning of each video, and another 1.8ml water was delivered following a 6-s blank period at the end of the video. The monkeys took 5-minute breaks between blocks (Figure S1A).

### Experimental stimuli

The stimuli used in this study were downloaded from YouTube. We applied Video Studio X8 (Corel Corporation, Canada) to edit these videos into 720P segments with 25 frames per second. In total, we prepared eighteen 30-s footage that were classified into three categories: 1) Primate content: with depiction of activities of monkeys; 2) Non-primate content: with activities of other species, including deer, lions, hippopotamus, hyenas, storks, rhinoceros, ostriches, penguins, and giraffes; 3) Scenery content: with depiction of dynamic naturalistic scenes without any animals.

### Ethogram analysis

Ethogram is used to describe a set of archetypal naturalistic behaviors by using descriptive terms and phrases of a species. For the collection of videos (see Stimuli), we constructed an inventory of behaviors by adapting the ethogram framework in Adams et al. (Adams et al., 2021), which is by far the most comprehensive ethogram analysis. The most common animal behaviors and their definitions presented in the videos were summarized in Table S2 (but note also (Partan, 2002)). Each video contains only a subset of ethogram features and no single video contains the full 52-item list (see Table S3 for description of the ethogram). Different elements of the ethogram were presented across different videos and across days. We did not fully control for the presence of all high-level features in the videos in this study. In contrast, by default, the low-level visual features exist in all videos. The behaviors for each video was manually registered by a custom program, the Tinbergen Alpha (Adams, 2014), producing a complete binary time-series of observable events.

### Low-level features extraction

Measures of video low-level features were extracted by applying Python and MATLAB for further modeling (Figure S2A-C). OpenCV package (Bradski & Kaehler, 2000) was called in Python for the calculation of luminance, contrast, and saturation. Luminance of each video frame was the mean of the pixel-wise luminosity, which was computed with the following equation l_pixel_ = 0.299 * R + 0.587 * G + 0.114 * B (Jack, 2008). Contrast of each frame was the standard deviation of the pixel-wise intensity distribution of the grayscale frame (Perfetto et al., 2020). Saturation was the mean pixel-wise S value of HSV color space that transformed from RGB color space (Jack, 2008). Motion was evaluated by the mean velocity magnitudes of optical flow by using the in-built Horn-Schunk algorithm in MATLAB (Bartels et al., 2008; Sliwa & Freiwald, 2017).

### Eye tracker and eye movement experiment

An infrared EyeLink 1000 Plus acquisition device (SR Research Ltd, Ottawa, CA) was used to track eye positions at a sampling rate of 1000 Hz. The Illuminator Module and the camera were positioned above the monkeys’ head. An angled infrared mirror was used to capture and re-coordinate monkeys’ eye positions. Each trial was initiated with a self-paced 2 s (range from 1.85 to 2.15 s) fixation on the white dot (50 × 50 pixels) centered on the screen.

### Electrophysiological recording and spike sorting

By using chronically implanted glass-coated electrodes from the right hemisphere (SC32, Gray Matter Research, LLC, USA) on monkeys Jupiter and Mercury, and by using single-wire tungsten microelectrode with 24 probes (LMA single shank, Microprobes, USA) on monkey Galen, we recorded multi-unit activities. In each recording session, the monkeys sat in chairs with their heads fixed. Headstage of multi-channel utility was connected to the SmartBox (NeuroNexus technologies, Inc., USA) acquisition system via an amplifier Intan adapter (RHD2000, Intan Technologies, USA) with 32 unipolar inputs. Microelectrodes impedance of each channel was in the range of 0.5 to 2.5 MΩ and measured at the beginning of the session. Spike waveforms above a set threshold were identified with a 1000 Hz online high-pass filter. Electrophysiological data collection was band-pass filtered from 0.1 to 5500 Hz and digitized at 30 kHz. Electrophysiological data from different sessions were treated as separate ones. Single units and their spikes were then identified based on peak amplitude, principal component, auto-correlation, and spike width by using Offline Sorter (Plexon Inc., USA). Units with an overall mean firing rate of fewer than 1 Hz across video presentations were excluded from further analysis.

Before recording, for monkeys Jupiter and Mercury, channels without spikes were manually advanced anti-clockwise to detect promised spike waveform. On any given day, the individual electrode was advanced at most 8 rounds (1 mm) with the step of one-eighth to 1 round (15.625 to 125 μm). For monkey G, a custom-designed recording grid (Delrin, 56 mm × 33.5 mm, 5 mm in thickness) with interlaced holes (0.8 mm in diameter and 0.8 mm apart from each other) was fixed on the plastic chips in the head-post. Then, an accommodated guide tube leads the 24 probes tungsten microelectrode (LMA single shank, Microprobes, USA), with 0.1 mm spacing between adjacent probes, through the skull and dura. A hydraulic microdrive (FHC Inc., USA) was used to drive the microelectrode into the target cortex, which was determined by MRI T1 images. By the end of the study, Jupiter and Mercury were scanned with CT. The location of each electrode was confirmed by mapping the CT image to MRI T1 images (Figure S1B). Histological recording sites were reconstructed based on the penetration depth of each electrode with the chamber coordinates and angles to the transverse plane.

### Surgical procedure for head-post and electrodes implantation

For monkeys Jupiter and Mercury, the surgeries consisted of 2 stages: head-post installation and electrodes implantation. Each stage was followed by a recovery period during which one dose of analgesic (Tolfedine, Vetoquinol, New Zealand) and antibiotics (Baytril, Bayer HealthCare LLC., Germany) were daily given via intramuscular injection according to body weight for one week. All medical operations and health care pre-post surgeries comply with the Institutional Animal Care and Use Committee guidelines at East China Normal University.

#### Head-post installation

Food and water were limited to 12 h before surgery. Forty-five minutes before the surgery, one dose of atropine sulfate (Shanghai Pharma, Changzhou Pharmaceutical Factory Co., Ltd., China) was injected to reduce saliva secretion during operations. Ten minutes later, one dose of Zoletil (Zoletil, Virbac, New Zealand) was injected for anesthetization before monkeys were transferred to the preparation room for shaving the head skin. Once the skin was prepared, the monkeys were placed on the stereotaxic apparatus mounted on the operating table. A mixture of oxygen and isoflurane was inhaled with the help of a ventilator. Dexamethasone (0.5mg/kg, Jilin Huamu Animal Health Products CO., LTD, China) was administered via intravenous transfusion with a 5% glucose-saline (Sake-biotech, China) injection at the beginning of the surgery to reduce the intracranial pressure and avoid bone inflammation during or after the surgery. Respiration, heart rate, blood pressure, expired CO_2_, and oxygen saturation were monitored during the whole surgical procedure. Body temperature was sustained at 37℃ with a constant temperature heater under the operating table. After successfully opening the epidermis and removing the subcutaneous tissues, an MRI-compatible polyether-ether-ketone (PEEK, Gray-Matter Research, Bozeman, USA) head post was cemented by acrylate cement (Refine Bright, Yamahachi Dental MFG.CO., Japan) which was then anchored with ceramic bone screws (Gray-Matter Research, Bozeman, USA) distributed on the anterior part of the skull. Sterilized saline was dropped to cool the hardened cement rapidly and clean the smoothed crumbs around the wound. Analgesics and anti-inflammatory were injected as required, when the ventilator was turned off and intravenous injection withdrew. MRI anatomical scans were acquired 4 months afterwards to aid subsequent implantation of the recording chambers.

#### Recording chamber implantation

Preoperative preparations were identical to the first stage. After opening the epidermis and removing the subcutaneous tissues, a craniotomy (5/8 inch in diameter) was manually drilled over the right hemisphere of the monkey, while the center of the chamber was pre-determined by the simulation of 3D Slicer (Kikinis et al., 2014). Next, the surface around the craniotomy was polished into a plane, and the medial wall was smoothed to only accommodate the chamber of the acquisition system. Then, 12 ceramic screws were placed for chamber fixation, and 2 stainless steel screws were anchored for grounding purposes. After that, the surrounding areas of the screws were tightly sealed with Super Bond (Sun Medical Co., Ltd., Japan), and the chamber was fixed with Palacos (Heraeus Medical, US) and acrylate cement.

Immediately after that, the monkey was transferred into an MRI scanner and imaged with a fiducial filled with gadopentetate dimeglumine (Shanghai Xudong Haipu Pharmaceutical Co., Ltd, China) that diluted 750 times. The center of the chamber was re-coordinated based on the modeling with fiducial marker by using 3D Slicer (monkey Jupiter: anterior-posterior (AP): -16.4 mm, medial-lateral (ML): 5.8 mm lateral to medial, 28°angle to the right and 14°angle to the posterior of the transverse plane; monkey Mercury: AP: -15.422 mm, ML: 7.549 mm, 25°angle to the right and 9.1° angle to the posterior; Figure S1B & 1C), covering paracentral of area 7a. Twenty-four hours later, the asepsis electrode-set was fitted into the chamber when monkeys were awake, and 44 rounds (5.5 mm) of each single electrode were gradually lowered anti-clockwise to penetrate through the dura and pia.

For monkey Galen, T1 images were scanned before surgery. After exposure of the skull and removal of the hypodermis, a lightweight acrylic cap was anchored by six titanium screws with acrylate cement for head-fixation. The cavity of the chamber was filled with a layer of cement, and two custom-designed plastic chips (10 mm ×56 mm, 5 mm in thickness) were stabilized over the hardened cement for recording grid restriction. At the end of the surgery, an acrylic resin cap was covered over the chamber. The monkey was allowed to rest for recovery.

## QUANTIFICATION AND STATISTICAL ANALYSIS

Data analysis was performed using custom software written in R, Python, and MATLAB.

### Feature selection with LASSO regression

We employed the ‘glmnet’ package (Friedman et al., 2017) in R language to build a linear model with elastic net regulation LASSO (least absolute shrinkage and selection operator) feature selection algorithm (Tibshirani, 1996). In contrast to the commonly used general linear model, LASSO regression has advantages for the present study. Annotation of ethograms produced a schematic binary time-series with 52 dimensions embedded within all the videos. An important feature of the ethogram is that some items are linearly correlated and not mutually exclusive. This implies that we might not be able to disentangle the internal structures/relationships among 52 ethogram items. For example, if the count of animals is equal or more than 2, the animal count must be larger than 1. A large number of regressors and multi-collinearity of simultaneous happenings tend to cause overfitting, which will increase the value of cost function, and reduce the explanatory power of the model. The LASSO regression will scale all variables, and shrink coefficients of less important predictors to 0 to filter out these redundant items from the model. In short, the LASSO algorithm selects features with non-zero coefficients by minimizing the prediction error of the model (Muthukrishnan & Rohini, 2016; Tibshirani, 1996; Zou & Hastie, 2005), which allowed us to determine which selected feature modulates the neural activity.

For each neuron, we concatenated the sequences of average superposed spike counts in 40 ms time-bin over 30 repetitions, as well as the time-series of ethograms, in the order of Non-primate, Primate, and Scenery video. A LASSO regression was constructed to model the modulation of 52 ethogram items and 4 low level features as variables on neuronal activity. With an increasing parameter λ, the algorithm iteratively penalized coefficients of all items gradually shrinking to zero (Figure S4A and S4B). An optimal λ was obtained by an in-built cross-validation procedure when the LASSO algorithm reached a minimum residual sum of squares. With the optimal λ parameter, these features with non-zero coefficients were then selected into the model (Figure S4C), implying that neural activity was effectively modulated by these selected features. For model validation, we performed a cross-validation with a model with features using random 80% of the sample data to predict the remaining 20% of the data (Figure S4D).

### Eye movement analysis

Raw eye movement data was converted from *.edf* format to *.asc* format. The following analyses were conducted by custom-coded R programs.

#### Saccade identification and saccadic modeling

We defined the duration of a saccadic event as the time elapsed from when the eye velocity exceeds 15° s^-1^ to when it returns below it (Goffart et al., 2017). In the present study, saccades were identified by the EyeLink 1000 Plus acquisition system with a ‘SACC’ marker during recording. In total, 41,646 saccadic behaviors were found with a mean amplitude of 11.20°with a mean saccadic duration of 70.35 ms.

For each trial, the eye movement timeline was aligned with the onset of movie stimulus. Then, the start moment and end moment of each saccade were divided by 0.04 s (40 ms, duration of one frame) to localize the start-frame and end-frame of that saccade. If the start-point and end-point were localized in one same frame, the corresponding frame would be valued with 1. If the start-point and end-point were localized in separate frames, the frames from the beginning of the saccade to the end of the saccade would all be valued with 1. The remaining frames without saccade were labeled with 0. By using this method, a frame-by-frame time serial binary saccadic event was generated trial-by-trial, and defined as the 57^th^ ethogram item. Then, a LASSO feature selection algorithm was performed for each neuron from monkey Galen to fit the neural activities with the binary labeling of the 52 ethogram-items, 4 low-level visual features, and the saccadic item.

#### Grid segregation and scan-path similarity

Grid segregation serves the purpose of dimensionality reduction of the data. The recorded x and y coordinates of fixations and saccades are mapped to a sequence by the formula:

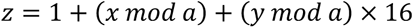

where the size of grids by the formula:

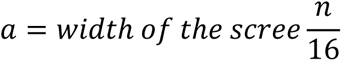

For monkey Galen, the screen is of 1440 ×900 and ⍺ = 90 in pixel. For monkeys P and K, the screen is of 2560 × 1440 and ⍺ = 160 in pixel. Irrespective of the screen dimension, the mapping creates a sequence that contains numbers ranging from 1 to 144. Each number represents the location of one corresponding fixation or saccade. For example, a fixation in Grid 1 is located at the bottom left corner of the screen, while a fixation in Grid 16 is located at the bottom right corner. Grid 17 is in the same column but one row above that of Grid 1.

Next, we calculated the x and y coordinates for the averaged eye position in each frame. The scan-path is defined by mapping the coordinates into grids to produce frame-by-frame eye position trajectories throughout the viewing of videos. The coordinates during blinks were filled with a linear interpolation by using 100 ms coordinates before and after a blink as the baseline. The scan-path similarity over viewings was estimated using pairwise correlation. To compare the variation in scan-path over repetitive viewings, correlation coefficient *r* was translated into *z* using a Fisher z transformation algorithm (Silver & Dunlap, 1987).

### Decoding analysis

We used a support vector machine (SVM) decoding method in the present study.

#### Video content type discriminability

A support vector machine (SVM) classifier with a leave-one-out cross-validation approach was performed on trial spike counts sequence binned in 1-s bins to quantify the representations of categorical natural episodes within neuronal activity with the help of the ‘e1071’ (Meyer et al., 2019) package in R. For each neuron, a multi-class decoder was trained on 87 trials (29 trials of each video), and tested on the remaining 3 trials using a one-versus-all method. Overall decoding performance was estimated as the average accuracy over 30 repetitions. At the same time, the decoding accuracy of each video was taken as the percentage of trials correctly predicted. To test the significance of decoding ability, we trained a multi-class classifier using spike sequences with randomly shuffled labels of training data and tested the left shuffled-label trials over 1000 repetitions. The statistical significance of real decoding performance was determined in comparison to the 95% percentile of shuffled decoding accuracy.

#### Temporal accumulation

We iteratively trained the SVM decoder utilizing spike counts in 1 s time bins from the 1^st^ to the 30^th^ time bin for the temporal accumulation decoding analysis to differentiate the video contents. A leave-one-out cross-validation training-testing SVM decoding approach was implemented at each accumulation timepoint. A multi-class decoder was trained on 87 trials (29 trials of each video), and tested on the remaining 3 trials using a one-versus-all method for each neuron at each timepoint. Spike count in the 1^st^ 1 s time-bin, for example, was used for the 1^st^ accumulation time point. The spike count sequences from the 1^st^ to 2^nd^ time bins were used in the decoding technique for the 2^nd^ accumulation timepoint, while sequences of spike count from the 1^st^, 2^nd^, and 3^rd^ time bins were used in the decoding approach for the 3^rd^ time point, and so on. We performed a similar decoding study for all individual timepoints to confirm that the accumulating effect is an intrinsic neural function rather than a momentary response activity (spiking in 1-s time bins) to stimuli (Figure S7). For the estimation of statistical significance, a similar permutation SVM decoding technique was used for corresponding accumulation and individual timepoints, respectively.

We defined three criteria to verify the accumulation neuron; (1) decoding accuracy in real firing sequences is significantly higher than corresponding label-shuffled firing sequences; (2) decoding accuracy in real firing sequences is significantly higher than corresponding individual timepoint decoding performance; (3) decoding performances in real firing sequences as a function of accumulated time point.

### Chi-squared simulation

A Chi-squared simulation procedure is used to determine the chance level of percentage of units modulated by specific selected features. The chance level is determined at 7.2% (or 27 /375 neurons, see blue dashed line in Figure 2B).

## SUPPLEMENTAL INFORMATION

Supplemental information can be found online.

## ACKNOWLEDGMENTS

This work received support from National Natural Science Foundation of China (32071060), Jiangsu Provincial Department of Science and Technology (BK20221267), and internal funding from School of Psychology and Cognitive Science (East China Normal University). We thank Emiliano Macaluso and Guohua Xu for their helpful comments on the manuscript.

## AUTHOR CONTRIBUTIONS

Conceptualization, L.W., S.C.K.; methodology, L.W., M.K., S.C.K.; investigation, L.W., X.Z., J.Y., M.C., F.Z., S.Z.; formal analysis, L.W., X.Z., J.Y., M.C.; visualization, L.W.; writing – original draft, L.W.; writing – review & editing, L.W., M.K., Y-D.Z., H.M.W., A.C., S.C.K.; supervision, H.M.W., S.C.K.; funding acquisition, S.C.K..

## Reference

Adams, G. (2014). Tinbergen Alpha. Release 1. http://dx.doi.org/10.5281/zenodo.13009 http://zenodo.org/record/13009#.VNj3Ji7w-7A.

Adams, G. K., Ong, W. S., Pearson, J. M., Watson, K. K., & Platt, M. L. (2021). Neurons in primate prefrontal cortex signal valuable social information during natural viewing. Philosophical Transactions of the Royal Society B, 376(1819), 20190666.

Andersen, R., Essick, G., & Siegel, R. (1987). Neurons of area 7 activated by both visual stimuli and oculomotor behavior. Experimental Brain Research, 67(2), 316–322.

Andersen, R. A., Bracewell, R. M., Barash, S., Gnadt, J. W., & Fogassi, L. (1990). Eye position effects on visual, memory, and saccade-related activity in areas LIP and 7a of macaque. Journal of Neuroscience, 10(4), 1176–1196.

Andric, M., Goldin-Meadow, S., Small, S. L., & Hasson, U. (2016). Repeated movie viewings produce similar local activity patterns but different network configurations. Neuroimage, 142, 613–627.

Baraduc, P., Duhamel, J.-R., & Wirth, S. (2019). Schema cells in the macaque hippocampus. Science, 363(6427), 635-639.

Barash, S., Bracewell, R. M., Fogassi, L., Gnadt, J. W., & Andersen, R. A. (1991). Saccade-related activity in the lateral intraparietal area. I. Temporal properties; comparison with area 7a. Journal of Neurophysiology, 66(3), 1095–1108.

Bartels, A., Zeki, S., & Logothetis, N. K. (2008). Natural vision reveals regional specialization to local motion and to contrast-invariant, global flow in the human brain. Cerebral cortex, 18(3), 705–717.

Bisley, J. W., Krishna, B. S., & Goldberg, M. E. (2004). A rapid and precise on-response in posterior parietal cortex. Journal of Neuroscience, 24(8), 1833–1838.

Bradski, G., & Kaehler, A. (2000). OpenCV. Dr. Dobb’s journal of software tools, 3.

Brodt, S., Gais, S., Beck, J., Erb, M., Scheffler, K., & Schönauer, M. (2018). Fast track to the neocortex: A memory engram in the posterior parietal cortex. Science, 362(6418), 1045–1048.

Brodt, S., Pöhlchen, D., Flanagin, V. L., Glasauer, S., Gais, S., & Schönauer, M. (2016). Rapid and independent memory formation in the parietal cortex. Proceedings of the National Academy of Sciences, 113(46), 13251–13256.

Bushnell, M. C., Goldberg, M. E., & Robinson, D. L. (1981). Behavioral enhancement of visual responses in monkey cerebral cortex. I. Modulation in posterior parietal cortex related to selective visual attention. Journal of Neurophysiology, 46(4), 755–772.

Cavanna, A. E., & Trimble, M. R. (2006). The precuneus: a review of its functional anatomy and behavioural correlates. Brain, 129(3), 564–583.

Clewett, D., DuBrow, S., & Davachi, L. (2019). Transcending time in the brain: How event memories are constructed from experience. Hippocampus, 29(3), 162–183.

Diomedi, S., Vaccari, F. E., Filippini, M., Fattori, P., & Galletti, C. (2020). Mixed selectivity in macaque medial parietal cortex during eye-hand reaching. Iscience, 23(10).

Erlich, J. C., Brunton, B. W., Duan, C. A., Hanks, T. D., & Brody, C. D. (2015). Distinct effects of prefrontal and parietal cortex inactivations on an accumulation of evidence task in the rat. Elife, 4, e05457.

Evangeliou, M. N., Raos, V., Galletti, C., & Savaki, H. E. (2009). Functional imaging of the parietal cortex during action execution and observation. Cerebral cortex, 19(3), 624–639.

Freedman, D. J., & Ibos, G. (2018). An integrative framework for sensory, motor, and cognitive functions of the posterior parietal cortex. Neuron, 97(6), 1219–1234.

Friedman, J., Hastie, T., Simon, N., Tibshirani, R., Hastie, M. T., & Matrix, D. (2017). Package ‘glmnet.’. Journal of Statistical Software, 33(1), 1–22.

Fusi, S., Miller, E. K., & Rigotti, M. (2016). Why neurons mix: high dimensionality for higher cognition. Current opinion in neurobiology, 37, 66–74.

Gardner, E. P., Babu, K. S., Reitzen, S. D., Ghosh, S., Brown, A. S., Chen, J., Hall, A. L., Herzlinger, M. D., Kohlenstein, J. B., & Ro, J. Y. (2007). Neurophysiology of prehension. I. Posterior parietal cortex and object-oriented hand behaviors. Journal of Neurophysiology, 97(1), 387–406.

Ghaem, O., Mellet, E., Crivello, F., Tzourio, N., Mazoyer, B., Berthoz, A., & Denis, M. (1997). Mental navigation along memorized routes activates the hippocampus, precuneus, and insula. Neuroreport, 8(3), 739–744.

Gilissen, S. R., & Arckens, L. (2021). Posterior parietal cortex contributions to cross-modal brain plasticity upon sensory loss. Current opinion in neurobiology, 67, 16–25.

Goffart, L., Bourrelly, C., & Quinet, J. (2017). Synchronizing the tracking eye movements with the motion of a visual target: Basic neural processes. Progress in Brain Research, 236, 243–268.

Gulli, R. A., Duong, L. R., Corrigan, B. W., Doucet, G., Williams, S., Fusi, S., & Martinez-Trujillo, J. C. (2020). Context-dependent representations of objects and space in the primate hippocampus during virtual navigation. Nature neuroscience, 23(1), 103–112.

Harvey, B. M., Klein, B. P., Petridou, N., & Dumoulin, S. O. (2013). Topographic representation of numerosity in the human parietal cortex. Science, 341(6150), 1123–1126.

Hasson, U., Chen, J., & Honey, C. J. (2015). Hierarchical process memory: memory as an integral component of information processing. Trends in cognitive sciences, 19(6), 304–313.

Hasson, U., Yang, E., Vallines, I., Heeger, D. J., & Rubin, N. (2008). A hierarchy of temporal receptive windows in human cortex. Journal of Neuroscience, 28(10), 2539–2550.

Honey, C. J., Thesen, T., Donner, T. H., Silbert, L. J., Carlson, C. E., Devinsky, O., Doyle, W. K., Rubin, N., Heeger, D. J., & Hasson, U. (2012). Slow cortical dynamics and the accumulation of information over long timescales. Neuron, 76(2), 423–434.

Jack, K. (2008). Color spaces. Digital Video and DSP, 15–29.

Jeunehomme, O., Heinen, R., Stawarczyk, D., Axmacher, N., & D’Argembeau, A. (2022). Representational dynamics of memories for real-life events. BioRxiv.

Jia, J., Puyang, Z., Wang, Q., Jin, X., & Chen, A. (2021). Dynamic encoding of saccade sequences in primate frontal eye field. The Journal of Physiology, 599(22), 5061–5084.

Johnston, W. J., Palmer, S. E., & Freedman, D. J. (2020). Nonlinear mixed selectivity supports reliable neural computation. PLOS computational biology, 16(2), e1007544.

Kano, F., & Hirata, S. (2015). Great apes make anticipatory looks based on long-term memory of single events. Current Biology, 25(19), 2513–2517.

Kikinis, R., Pieper, S. D., & Vosburgh, K. G. (2014). 3D Slicer: a platform for subject-specific image analysis, visualization, and clinical support. In Intraoperative imaging and image-guided therapy (pp. 277-289). Springer.

Klein, J. T., Deaner, R. O., & Platt, M. L. (2008). Neural correlates of social target value in macaque parietal cortex. Current Biology, 18(6), 419–424.

Kravitz, D. J., Saleem, K. S., Baker, C. I., & Mishkin, M. (2011). A new neural framework for visuospatial processing. Nature Reviews Neuroscience, 12(4), 217–230.

Lieberman, M. D. (2007). Social cognitive neuroscience: a review of core processes. Annu. Rev. Psychol., 58, 259–289.

Meyer, D., Dimitriadou, E., Hornik, K., Weingessel, A., Leisch, F., Chang, C.-C., Lin, C.-C., & Meyer, M. D. (2019). Package ‘e1071’. The R Journal.

Morecraft, R., Cipolloni, P., Stilwell-Morecraft, K., Gedney, M., & Pandya, D. (2004). Cytoarchitecture and cortical connections of the posterior cingulate and adjacent somatosensory fields in the rhesus monkey. Journal of Comparative Neurology, 469(1), 37–69.

Mosher, C. P., Zimmerman, P. E., & Gothard, K. M. (2014). Neurons in the monkey amygdala detect eye contact during naturalistic social interactions. Current Biology, 24(20), 2459–2464.

Murray, J. D., Bernacchia, A., Freedman, D. J., Romo, R., Wallis, J. D., Cai, X., Padoa-Schioppa, C., Pasternak, T., Seo, H., & Lee, D. (2014). A hierarchy of intrinsic timescales across primate cortex. Nature neuroscience, 17(12), 1661–1663.

Murray, J. D., Jaramillo, J., & Wang, X.-J. (2017). Working memory and decision-making in a frontoparietal circuit model. Journal of Neuroscience, 37(50), 12167–12186.

Muthukrishnan, R., & Rohini, R. (2016). LASSO: a feature selection technique in predictive modeling for machine learning. 2016 IEEE international conference on advances in computer applications (ICACA),

Partan, S. (2002). Single and multichannel signal composition: facial expressions and vocalizations of rhesus macaques (Macaca mulatta). Behaviour, 139(8), 993–1027.

Perfetto, S., Wilder, J., & Walther, D. B. (2020). Effects of spatial frequency filtering choices on the perception of filtered images. Vision, 4(2), 29.

Raffi, M., & Siegel, R. M. (2007). A functional architecture of optic flow in the inferior parietal lobule of the behaving monkey. PLoS one, 2(2), e200.

Ranganath, C., & Ritchey, M. (2012). Two cortical systems for memory-guided behaviour. Nature Reviews Neuroscience, 13(10), 713–726.

Reagh, Z. M., & Ranganath, C. (2021). A cortico-hippocampal scaffold for representing and recalling lifelike events. BioRxiv.

Rigotti, M., Barak, O., Warden, M. R., Wang, X.-J., Daw, N. D., Miller, E. K., & Fusi, S. (2013). The importance of mixed selectivity in complex cognitive tasks. Nature, 497(7451), 585–590.

Robinson, D. L., Goldberg, M. E., & Stanton, G. B. (1978). Parietal association cortex in the primate: sensory mechanisms and behavioral modulations. Journal of Neurophysiology, 41(4), 910–932.

Runyan, C. A., Piasini, E., Panzeri, S., & Harvey, C. D. (2017). Distinct timescales of population coding across cortex. Nature, 548(7665), 92–96.

Russ, B. E., & Leopold, D. A. (2015). Functional MRI mapping of dynamic visual features during natural viewing in the macaque. Neuroimage, 109, 84–94.

Satpute, A. B., & Lieberman, M. D. (2006). Integrating automatic and controlled processes into neurocognitive models of social cognition. Brain research, 1079(1), 86–97.

Silver, N. C., & Dunlap, W. P. (1987). Averaging correlation coefficients: should Fisher’s z transformation be used? Journal of applied psychology, 72(1), 146.

Sliwa, J., & Freiwald, W. A. (2017). A dedicated network for social interaction processing in the primate brain. Science, 356(6339), 745–749.

Snyder, L. H., Batista, A., & Andersen, R. A. (1997). Coding of intention in the posterior parietal cortex. Nature, 386(6621), 167–170.

Testard, C., Tremblay, S., & Platt, M. (2021). From the field to the lab and back: neuroethology of primate social behavior. Current opinion in neurobiology, 68, 76–83.

Tibshirani, R. (1996). Regression shrinkage and selection via the lasso. Journal of the Royal Statistical Society: Series B (Methodological), 58(1), 267–288.

Vaccari, F. E., Diomedi, S., Filippini, M., Hadjidimitrakis, K., & Fattori, P. (2022). New insights on single-neuron selectivity in the era of population-level approaches. Frontiers in Integrative Neuroscience, 16, 929052.

Wallach, A., Melanson, A., Longtin, A., & Maler, L. (2021). Mixed selectivity coding of sensory and motor social signals in the thalamus of a weakly electric fish. BioRxiv.

Wang, L., Zuo, S., Cai, Y., Zhang, B., Wang, H., & Kwok, S. C. (2020). Fallacious reversal of event-order during recall reveals memory reconstruction in rhesus monkeys. Behavioural Brain Research, 394, 112830.

Zhang, B., Wang, F., Zhang, Q., & Naya, Y. (2022). Distinct networks coupled with parietal cortex for spatial representations inside and outside the visual field. Neuroimage, 252, 119041.

Zou, H., & Hastie, T. (2005). Regularization and variable selection via the elastic net. Journal of the royal statistical society: series B (statistical methodology), 67(2), 301–320.

Zuo, S., Wang, L., Shin, J. H., Cai, Y., Zhang, B., Lee, S. W., Appiah, K., & Kwok, S. C. (2020). Behavioral evidence for memory replay of video episodes in the macaque. Elife, 9, e54519.

