## Supplementary information for "Mixed selectivity coding of content-temporal detail by dorsomedial posterior parietal neurons"

This manuscript contains four (4) supplemental tables, twelve (8) supplemental figures, and two (2) supplemental Movies.

Table S1. Lists of videos.

Table S2. Neural and eye-tracking data for each of the monkeys.

Table S3. Description and scoring of the ethogram.

Table S4. Full listing of the ethogram of the 18 videos.

Figure S1. Experimental procedure and recording sites. Related to Figure 1.

Figure S2. Demonstration of neural activities modulated by multiplex features embedded in an example video. Related to Figure 1 and Figure 2.

Figure S3. Low-level features extraction. Related to Figure 2.

Figure S4. Feature selection with LASSO. Related to Figure 2.

Figure S5. Neural modulations were not affected by neuronal transmission latency. Related to Figure 2.

Figure S6. Neural activity in dmPPC cannot be accounted for by saccadic eye movements. Related to Figure 2.

Figure S7. Illustration of how spikes were used for SVM training and testing for video content-type decoding for accumulation.

Figure S8. Information accumulation is not affected by the length of time-bin sequences. Related to Figure 5.

Other Supplemental Material for this manuscript includes the following:

Movie S1 (.mp4 format). Population of neurons responding to a Primate video.

Movie S2 (.mp4 format). Demonstration for dmPPC cell ensembles for their mixed selectivity coding.

Table S1 Lists of videos.

| Boundary | Content-type (Video's order) | | | List |
|---|---|---|---|---|
| | Primate | Non-primate | Scenery | |
| B0 | PB002.mp4 (1) | NB002.mp4 (2) | SB002.mp4 (3) | 1 |
| | PB003.mp4 (4) | NB003.mp4 (5) | SB003.mp4 (6) | 2 |
| B1 | PB102.mp4 (7) | NB102.mp4 (8) | SB102.mp4 (9) | 3 |
| | PB103.mp4 (10) | NB103.mp4 (11) | SB103.mp4 (12) | 4 |
| B2 | PB202.mp4 (13) | NB202.mp4 (14) | SB202.mp4 (15) | 5 |
| | PB203.mp4 (16) | NB203.mp4 (17) | SB203.mp4 (18) | 6 |

Note: For each video, the first letter stands for the video content-type (Primate/Non-Primate/Scenery), the next two strings represent boundary conditions (0/1/2), and the last two strings refer to an ordinal number of the video. For example, PB002.mp4 is a video containing primate content-type without boundaries. The numbers in brackets correspond to the ordinal order of videos shown in Figure S2B. The factor “boundary” was not considered in this current manuscript.

Table S2 Composition of neural and eye-tracking data for monkeys.

| Monkey | Jupiter | Mercury | Galen | K | P |
|---|---|---|---|---|---|
| Eye tracking | | | + | + | + |
| Electrophysiological recording | + | + | + | | |

Note: + indicates that data was acquired.

Table S3 Description and scoring of the ethogram.

| Observation | Levels | Description |
|---|---|---|
| camera movement | zooming in | the camera is holding a mostly stable image, but is zooming in |
| | zooming out | the camera is holding a mostly stable image, but is zooming out |
| | tracking | the camera is tracking the movements of a animal, holding it largely stable against the moving background |
| | panning | the camera is moving without tracking anything, and the scene is still |
| | slewing | the camera is moving wildly and the scene is difficult to view |
| face visibility | visible | faces are visible |
| | side face | at least one eye is visible |
| | direct face | two eyes are visible |
| | eye contact | as ‘direct face’, but the animal also appears to be looking directly at the camera |
| count more than | 1 | the number of animals visible in the scene, scored in approximately spaced levels |
| | 2 | |
| | 3 | |
| | 4 | |
| | 5 | |
| ano-genital area (AGA) visibility | none | the AGA of no animals is visible. |
| | visible (unsexed) | at least one animal’s AGA is visible, but none are prominent on the screen |
| | male | at least one female animal’s AGA is prominently visible |
| | female | at least one female animal’s AGA is prominently visible |
| | prominent | at least one animal’s AGA is prominently visible, but the observer is unable to sex the animal |
| forage | forage | one or two animals are in a foraging state, characterized by various specific actions such as manipulating or ingesting food |
| | group forage | three or more animals are engaged in foraging behaviors; due to the rapid increase in complexity of coding specific actions with this number of foraging animals, specific actions are only coded during the ‘forage’ condition |
| | drink | any visible animal is drinking water |
| | search | searching for a food item by manipulating foliage, ground substrate, or a pile of provisioned food items |
| | grasp food | reaching for a food item; begins at initiation of the reach movement and ends once the food item is in hand |
| | hold food | a food item is visibly held in hand or foot |
| | hold food in mouth | a food item is visibly held in the mouth; carrying food in cheek pouches is not include |
| | manipulate food | active manipulate of a food item other than grasping or ingesting, e.g. ‘washing’ |

(*continued*)

Table S3 (*continued*)

| Observation | Levels | Description |
|---|---|---|
| | ingest food | a food item is brought to the mouth: begins at initiation of arm movement and ends when the food item is consumed or the hand leaves the mouth area; includes movements of food item towards the mouth that do not actually result in consumption, e.g. 'sniffing' |
| | ingest grooming manipulate | a hand is brought from a grooming target (self or partner) to the mouth, as if the animal is consuming a parasite plucked from the skin |
| | chew | rhythmic jaw movements; an entire bout of chewing is coded rather than individual bites |
| | retrieve from pouch | a food item is brought out from the cheek pouches |
| groom | allogroom | grooming behavior, involve one animal uses its hands, mouth, or other part of its body to touch another animal with finely controlled hand and finger movements |
| | solicit allogroom | an animal approaches another animal and sits or lies down, presenting itself for grooming |
| | scratch | rhythmic, vigorous movement of the hand or foot against the animal's own body |
| | autogroom | self-directed grooming behavior |
| aggression | strike | a animal makes brief, aggressive physical contact with another |
| | grapple | two animals engage in prolonged aggressive physical contact |
| | lunge | an animal makes a short, aggressive movement toward another animal |
| | withdraw | an animal backs away from another animal while making aggressive or submissive displays |
| | charge | an animal makes a rapid, prolonged aggressive movement toward another animal |
| | chase | one animal charges another animal as it flees |
| | flee | an animal moves away from another animal at high speed |
| | threaten | an animal performs a threat display, characterized by a round open mouth, prolonged staring, head bobbing, piloerection, and erect posture |
| | mounted | one animal mounts another animal of its own species |
| | submit | an animal performs a submissive display, characterized by bearing teeth, squeaking and withdrawn posture |
| | displace | one animal walks directly toward or near another, which moves away |

(*continued*)

Table S3 (*continued*)

| Observation | Levels | Description |
|---|---|---|
| | lean away | an animal posturally shifts away from an approaching conspecific without fully withdrawing or displacing |
| | avoid | an animal pauses or alters course during movement to maintain greater distance to another animal |
| | branch display | an animal vigorously shakes a branch or branch-like object (e.g. a metal pole) |
| | none | no aggression is present in the scene |
| | unidirectional | a single individual is displaying aggressive behaviors |
| | bidirectional | two individuals are displaying aggressive behaviors toward each other |
| | joint | two or more individuals are jointly displaying aggressive behaviors, the target(s) of which may or may not be visible |
| | intercoalition | two ‘coalitions’ are displaying aggressive behaviors toward each other |

Table S4 Full listing of the ethogram of the 18 videos. Color in each cell denotes the presence of the associated feature (row) in the video (column). The three colors refer to the three different video content-types (green = Primate, blue = Non-primate, and red = Scenery).

| | PB002 | NB002 | SB002 | PB003 | NB003 | SB003 | PB102 | NB102 | SB102 | PB103 | NB103 | SB103 | PB202 | NB202 | SB202 | PB203 | NB203 | SB203 |
|---|---|---|---|---|---|---|---|---|---|---|---|---|---|---|---|---|---|---|
| luminance | ■ | ■ | ■ | ■ | ■ | ■ | ■ | ■ | ■ | ■ | ■ | ■ | ■ | ■ | ■ | ■ | ■ | ■ |
| contrast | ■ | ■ | ■ | ■ | ■ | ■ | ■ | ■ | ■ | ■ | ■ | ■ | ■ | ■ | ■ | ■ | ■ | ■ |
| saturation | ■ | ■ | ■ | ■ | ■ | ■ | ■ | ■ | ■ | ■ | ■ | ■ | ■ | ■ | ■ | ■ | ■ | ■ |
| optical flow | ■ | ■ | ■ | ■ | ■ | ■ | ■ | ■ | ■ | ■ | ■ | ■ | ■ | ■ | ■ | ■ | ■ | ■ |
| camera zooming in | ■ | | | | | | ■ | | | ■ | | | | | | | | |
| camera zooming out | ■ | | | ■ | | | | | | ■ | | | | | | | | |
| camera tracking | ■ | | ■ | ■ | ■ | ■ | ■ | ■ | ■ | ■ | ■ | ■ | | ■ | ■ | ■ | ■ | ■ |
| camera panning | ■ | | | ■ | | | ■ | | | ■ | ■ | | | | | | | |
| camera slewing | | | | | | | | | | | | | | | | | | |
| visible face | ■ | ■ | | ■ | ■ | | ■ | ■ | | ■ | ■ | | ■ | ■ | | ■ | ■ | |
| side face | ■ | ■ | | ■ | ■ | | ■ | ■ | | ■ | ■ | | ■ | ■ | | ■ | ■ | |
| direct face | ■ | ■ | | ■ | | | ■ | | | ■ | ■ | | | ■ | | ■ | ■ | |
| eye contact | ■ | ■ | | ■ | | | ■ | | | | | | | ■ | | ■ | | |
| count >= 1 | ■ | ■ | | ■ | ■ | | ■ | ■ | | ■ | ■ | | ■ | ■ | | ■ | ■ | |
| count >= 2 | ■ | ■ | | ■ | ■ | | ■ | ■ | | ■ | ■ | | ■ | ■ | | ■ | ■ | |
| count >= 3 | ■ | ■ | | ■ | | | ■ | ■ | | ■ | ■ | | | ■ | | ■ | ■ | |
| count >= 4 | ■ | ■ | | ■ | | | ■ | ■ | | ■ | | | | ■ | | ■ | ■ | |
| count >= 5 | | ■ | | | | | ■ | ■ | | | | | | ■ | | | ■ | |
| visible genitals | ■ | | | ■ | | | ■ | ■ | | ■ | | | | | | ■ | | |
| prominent genitals | ■ | | | ■ | | | ■ | | | ■ | | | | | | ■ | | |
| male genitals | ■ | | | ■ | | | ■ | | | ■ | | | | | | ■ | | |
| female genitals | | | | | | | | | | ■ | | | | | | ■ | | |
| foraging | | | | ■ | | | | | | ■ | ■ | | | | | | | |
| group foraging | | | | | | | | | | | ■ | | | | | | | |
| drink | | | | | | | | | | | | | | | | | | |
| search | ■ | | | | | | ■ | | | | ■ | | | | | | | |
| grasp food | | | | ■ | | | ■ | | | ■ | ■ | | | | | | | |
| hold food | | | | ■ | | | | | | ■ | | | | | | | | |
| hold food in mouth | | | | ■ | | | ■ | | | ■ | | | | | | | | |
| manipulate food | | | | ■ | | | | | | | | | | | | | | |
| ingest food | | | | ■ | | | ■ | | | ■ | | | | | | | | |
| ingest grooming manipuland | | | | | | | | | | | | | | | | | | |
| chew | | | | ■ | | | | | | ■ | | | | | | | | |
| retrieve from cheek pouch | | | | | | | | | | | | | | | | | | |
| allogroom | | | | | | | | | | ■ | | | | | | | | |
| solicit allogroom | | | | | | | | | | | | | | | | | | |
| scratch | ■ | | | | | | ■ | | | | | | | | | | | |
| autogroom | | | | | | | | | | | | | | | | | | |
| strike | | | | ■ | | | | | | ■ | | | | ■ | | ■ | | |
| grapple | | | | | | | | | | | | | | | | ■ | | |
| lunge | | | | ■ | | | | | | | | | | | | ■ | | |
| withdraw | | | | ■ | | | | | | | | | | | | | | |
| charge | | | | | | | | | | | | | | | | | | |
| chase | | | | | | | | | | ■ | | | ■ | ■ | | ■ | | |
| flee | | | | ■ | | | | | | | | | ■ | ■ | | | | |
| threaten | | | | | | | | | | | | | | | | | | |
| mounted threaten | | | | | | | | | | | | | | | | ■ | | |
| submit | | | | | | | | | | | | | | | | | | |
| displace | | | | | | | | | | | | | | | | | | |
| lean away | | | | | | | | | | | | | | | | | | |
| avoid | | | | | | | | | | | | | | | | | | |
| branch display | | | | | | | | | | | | | | | | | | |
| any aggression | | | | ■ | | | | | | ■ | | | ■ | ■ | | ■ | | |
| mutual aggression | | | | | | | | | | | | | | | | ■ | | |
| joint aggression | | | | | | | | | | | | | | | | | | |
| intercoalition aggression | | | | | | | | | | | | | | | | | | |

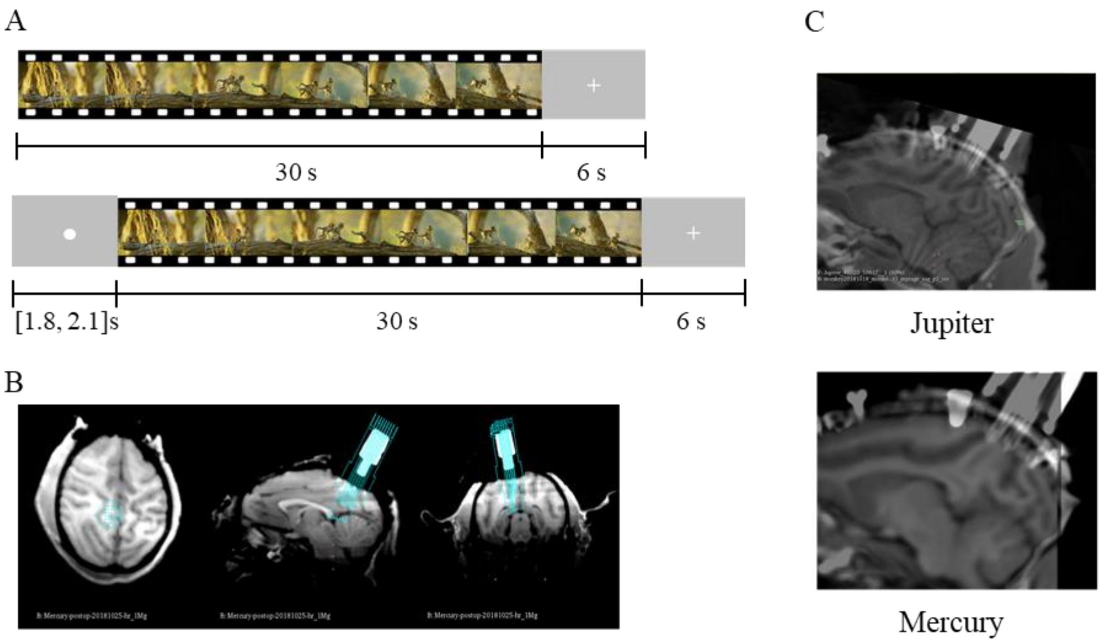


**Figure S1. Experimental procedure and recording sites**. (**A**) Each trial consists of 30-s video display and a 6-s reward period. In each block, 15 presentations (5 for each content-type) were presented in random order. There were 6 blocks in total. We added a self-paced fixation start (1.8 to 2.1 s) for monkeys Galen, K and P. (**B**) Simulated anatomical sites for the recording chamber shown in 3D Slicer (left), and with electrodes penetration trajectory (middle: sagittal plane; right: coronal plane), in cyan. White cylinder depicts the chamber filled with fiducial markers. (**C**) T1 MRI images overlaid on CT images indicating the actual position of electrodes in the dmPPC at the beginning of testing.

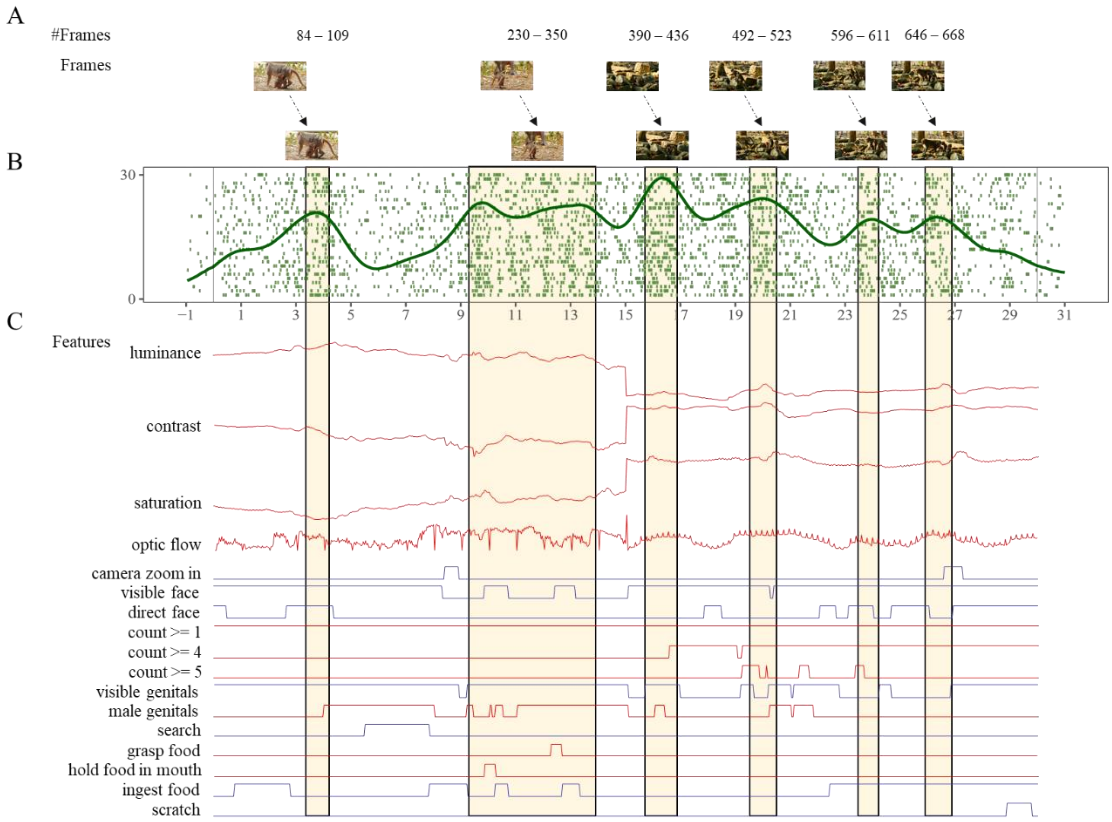


**Figure S2. Demonstration of neural activities modulated by multiplex features embedded in an example video.** (**A**) Segments of increased neural spiking with their corresponding frames. (**B**) Raster plot of an example neuron's (*#PC0056*) spiking in response to a Primate video (PB102). (**C**) Features selected by LASSO algorithms modulate the neural activities. Lines stand for the feature values of low-level visual features and ethogram variables across the video. Red and blue denote the positive and negative modulation of features, respectively. Related to Figure 1 and Figure 2.

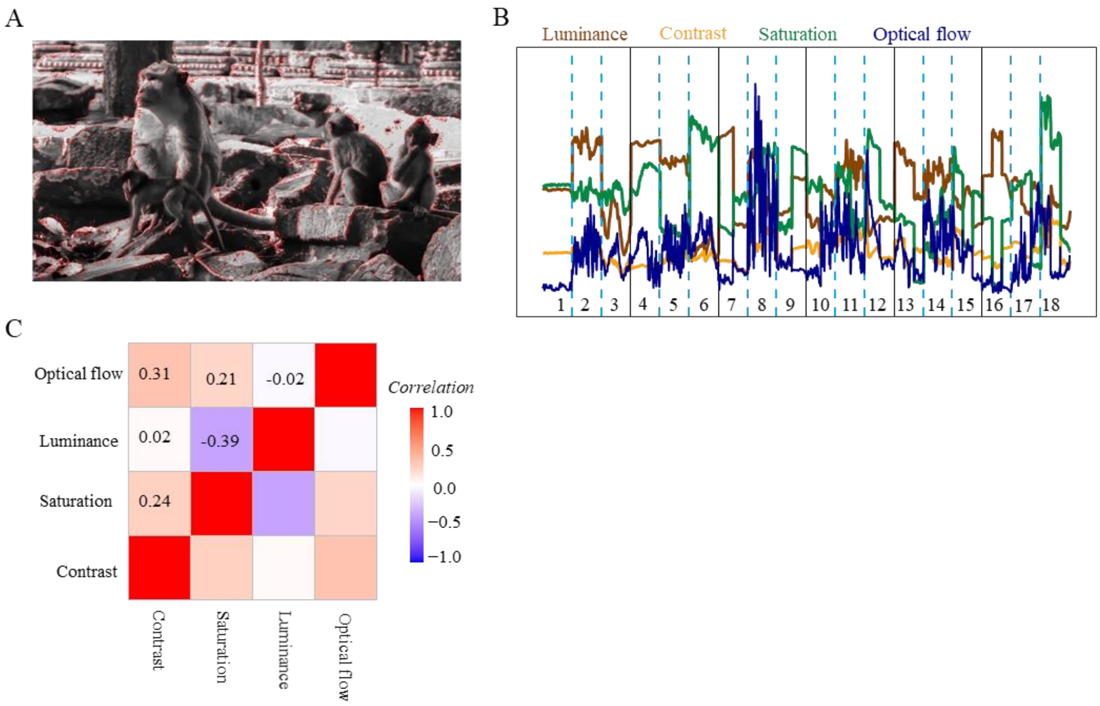


**Figure S3. Low-level features extraction.** (**A**) Example of optical flow of one movie frame shown in red vector arrows with amplitude and directions. (**B**) Time course of 4 low-level features. The numbers indicate the videos, with the 18 videos listed in the same order in Table S4. (**C**) The pair-wised correlation for the 4 low-level features across time for all 18 videos.

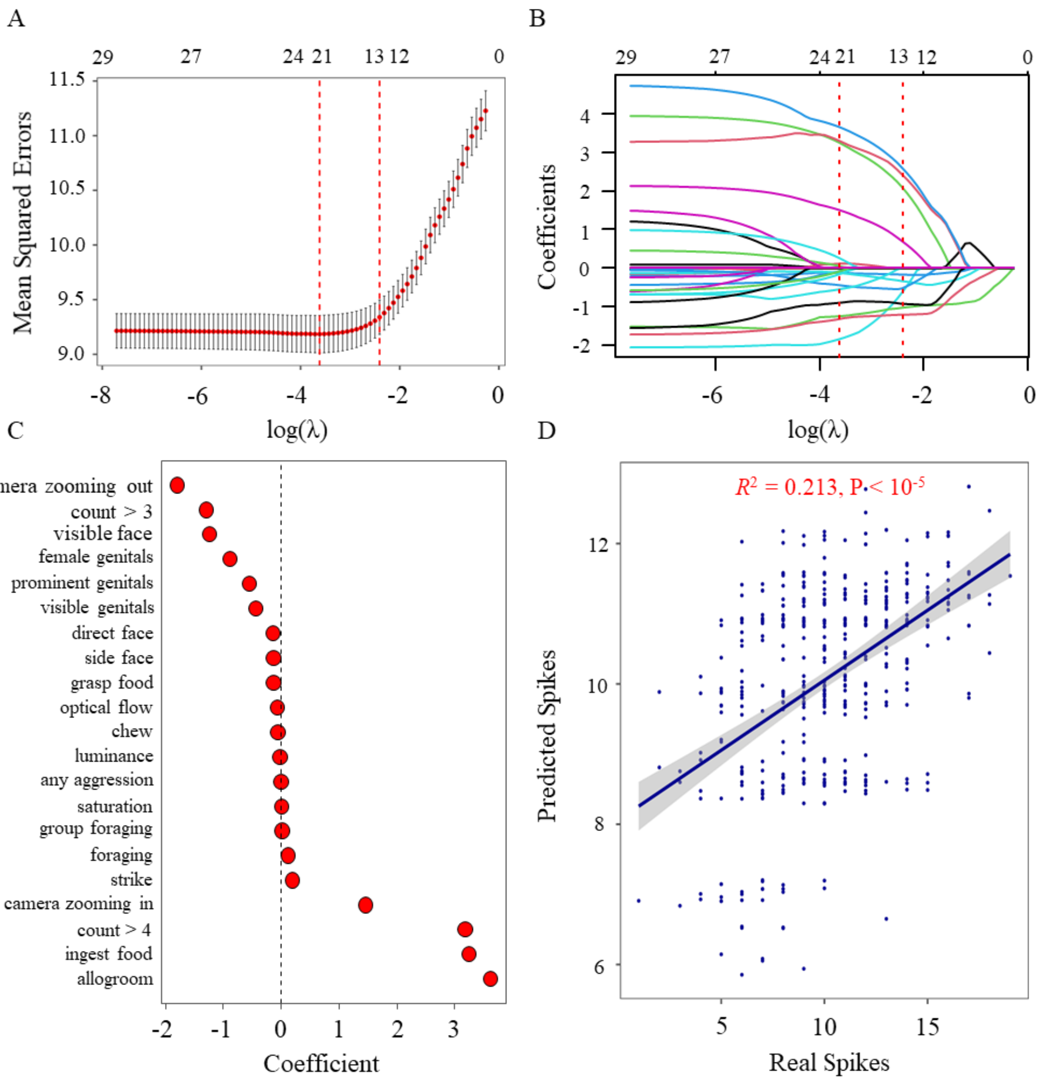


**Figure S4. Feature selection with LASSO.** (**A, B**) We fitted a least absolute shrinkage and selection operator regression model (LASSO) with spike counts in 40-ms time bin as dependent variable and 52 ethogram items and 4 low-level features as regressors. The algorithm punishes the coefficient of less important variables to zero along with the gradual increase of parameter log(λ). A variable would be filtered out of the model when its coefficient is punished to 0. A 10-fold cross-validation procedure was used to determine the value of lambda as the model produced minimal mean squared errors (MSE). For an example neuron #PC0087, the algorithm yielded an optimal model with 21 non-zero coefficient variables when log(λ) = -3.43. The red dashed lines represent the largest λ where the MSE is within one standard error of the minimal MSE. Solid curves in B refer to the coefficient path of variables. The numbers on top indicate the number of non-zero coefficient variables in the optimal model. **(C)** A set of non-zero coefficient variables produced by the model at minimal MSE. (**D**) For validation of the optimal model, a regression model, built with 80% training dataset to test 20% testing dataset, showed a significant prediction ability ($F(1, 450) = 121.5$, $R^2 = 0.213$, slope = 0.200).

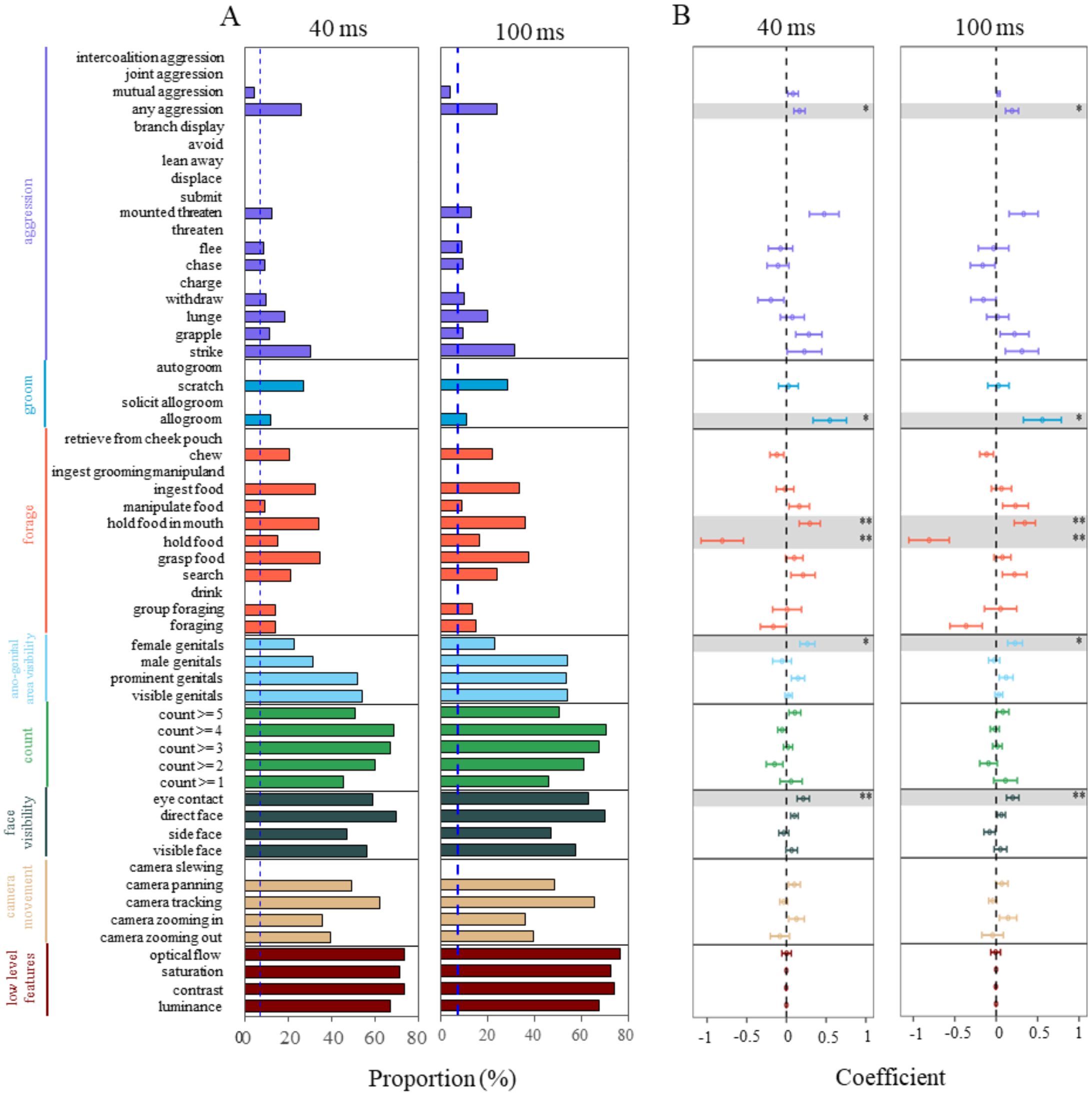


**Figure S5. Neural modulations were not affected by neuronal transmission latency.** Both the proportion (A) and coefficients (B) of the LASSO-selected features exhibited no statistically significant changes after 40-ms and 100-ms latency realignments in comparison to using 0-ms latency alignment. Error bars: SEM across neurons. * P < 0.05, ** P < 0.01. Related to Figure 2.

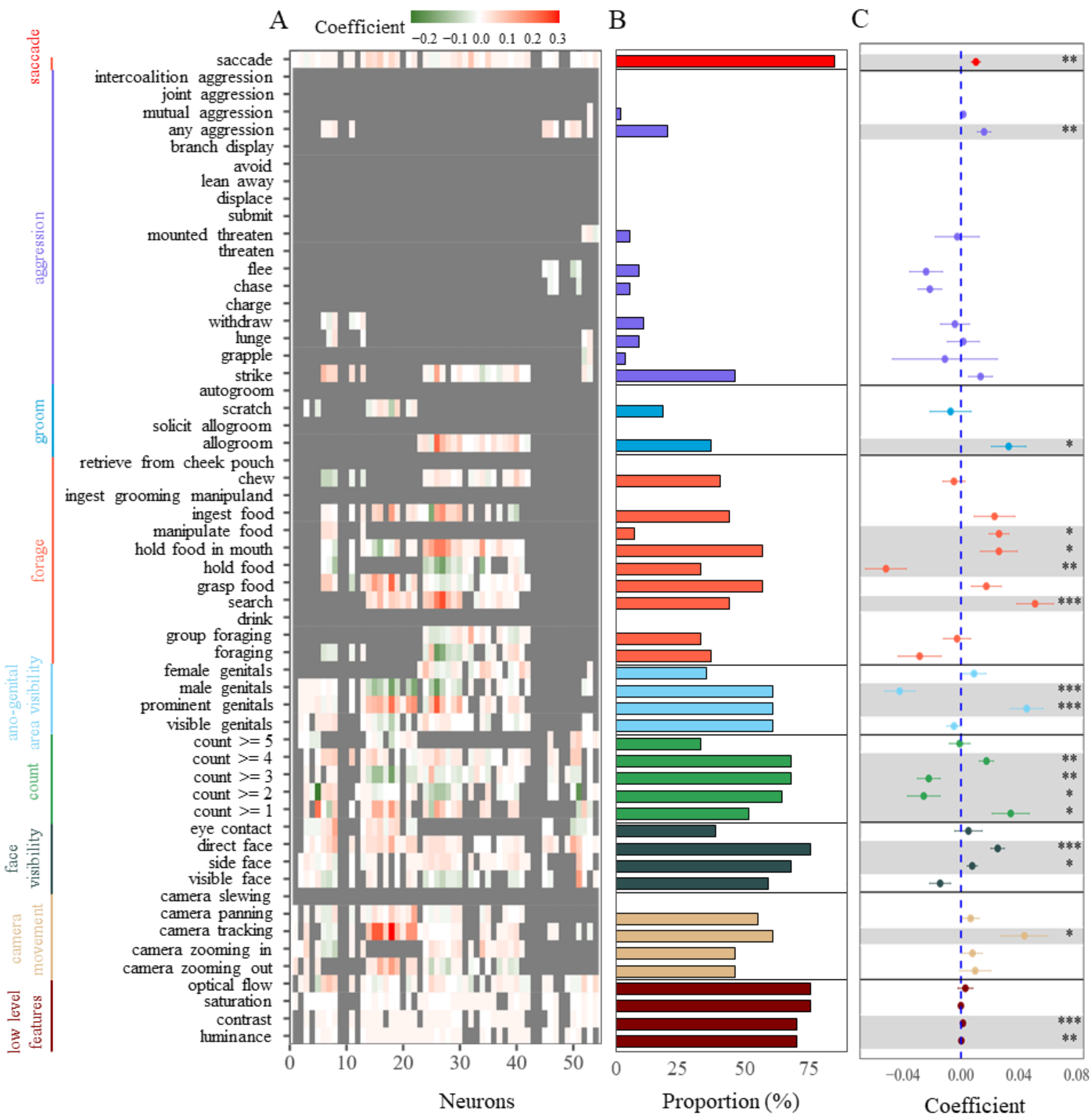


**Figure S6. Neural activity in dmPPC cannot be accounted for by saccadic eye movements.** For Monkey Galen eye movements were monitored with EyeLink 1000 and neural activities were recorded simultaneously during free viewing, we identified trial by trial saccadic eye movements and set saccadic as the 57$^{th}$ feature to evaluate the neural modulation by eye movement. (A) Effects of neuronal responses to saccadic eye movements and 56-item ethograms obtained from LASSO regression trial-by-trial. (B) Proportion of neurons responsive to each item. (C) LASSO coefficient for each item tested against zero. The proportion and coefficient of neurons responding to each of the 56 key items showed no statistical difference from those when no saccadic data were included as a feature. Error bars: SEM across neurons. * $P < 0.05$, ** $P < 0.01$, *** $P < 0.001$. Related to Figure 2.

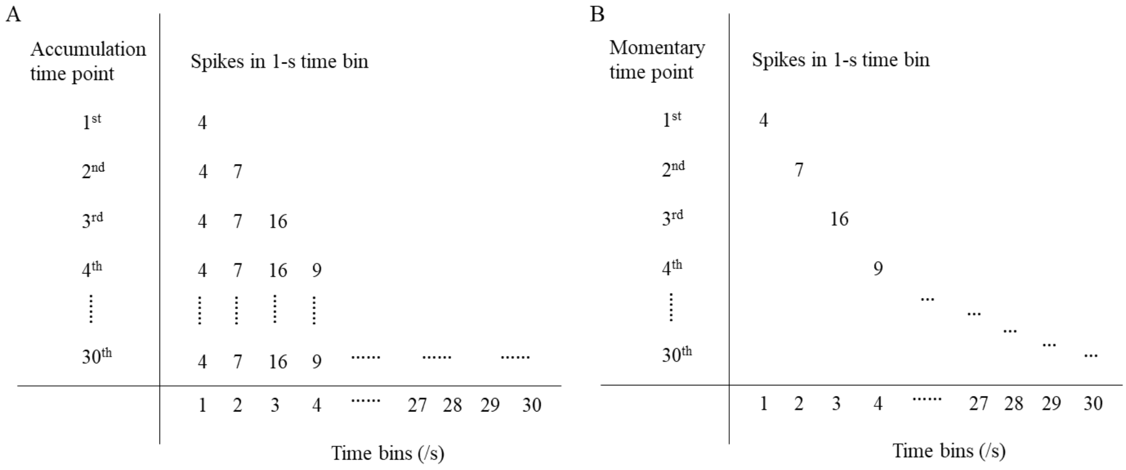


**Figure S7.** Illustration of how spikes were used for SVM training and testing for video content-type decoding for accumulation (A) and momentary conditions (B), respectively.

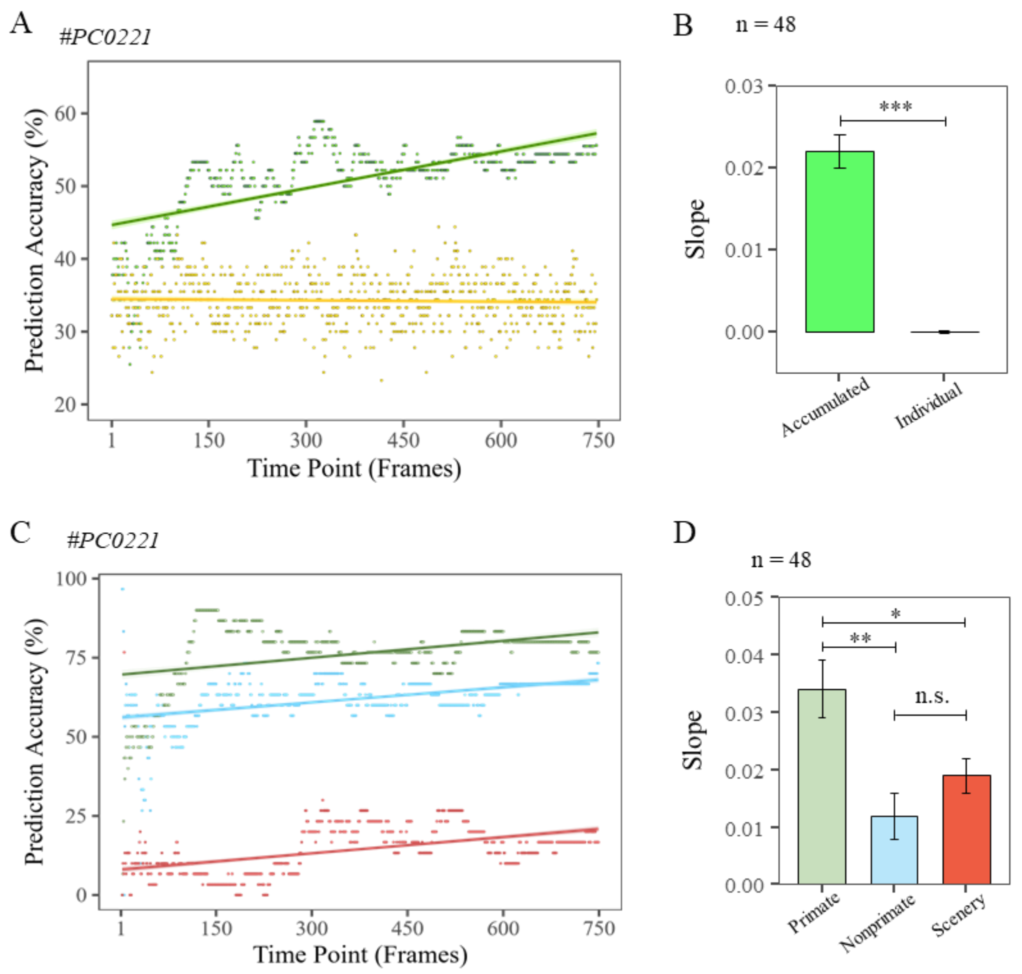


**Figure S8. Information accumulation is not affected by the length of time-bin sequences.** (A) In this set of analysis, we repeated the accumulation decoding analysis with cumulative spikes in 40-ms time bins from the 1st to 750th frame (green dots and line) and compared it to using neural activity in each individual frame duration (yellow dots and line in panel A). The results showed that the decoding performance of the example neuron (#PC0221) maintained its positive correlation with cumulative spiking sequences ($R^2$ = 0.433, $P < 10^{-5}$) but not with momentary neural activity ($R^2$ = 0.001, P = 0.294). **(B)** T-test showed that the mean slope of using cumulative spiking sequences (n = 48) was stronger than using momentary 40-ms binned decoding performance on a population level (t(47) = 10.947, P < 0.001, Cohen's D = 1.580), replicating our main results in Figure 5. **(C)** The decoding performance for each video content-type as a function of cumulative spiking sequences with 40-ms time bins (Primate: $R^2$ = 0.138, $P < 10^{-5}$; Non-primate: $R^2$ = 0.213, $P < 10^{-5}$; Scenery: $R^2$ = 0.236, $P < 10^{-5}$). (D) One-way ANOVA test and post hoc analysis revealed that neurons in dmPPC had faster accumulation speed and stronger magnitude for primate video content-type than the other two content-types (F(2, 94) = 6.753, P = 0.002, $\eta^2$ = 0.126; $t_{\text{Primate – Non-primate}}$ = 3.610, P = 0.001, Cohen's D = 0.782; $t_{\text{Primate – Scenery}}$ = 2.400, P = 0.037, Cohen's D = 0.520). Related to Figure 5.
